# Endothelial protein C receptor CD201 is a better marker than SCA1 to identify mouse long-term reconstituting hematopoietic stem cells following septic challenge

**DOI:** 10.1101/2025.11.13.688157

**Authors:** Kavita Bisht, Valérie Barbier, Svetlana Shatunova, Ingrid G. Winkler, Jean-Pierre Lévesque

**Affiliations:** Mater Research Institute, The University of Queensland, Woolloongabba, Queensland, Australia

## Abstract

Stem cell antigen-1 (SCA1) is widely used to identify mouse hematopoietic stem cells (HSC) and multipotent progenitors (MPP) among lineage-negative KIT^+^ (LK) cells. However, SCA1 is expressed only in a few inbred mouse strains and becomes strongly upregulated on LK cells following in vivo challenge with interferons, lipopolysaccharide (LPS) or pathogens leading to incorrect analysis of HSC function subsets and delineation of HSC, MPP and lineage-restricted progenitor subsets. Endothelial protein C receptor CD201can be used as an alternative marker for mouse and even human HSC. However, whether CD201 expression changes following infectious challenge is unknown. Unlike SCA1, CD201 expression did not change on mouse LK cells in response to LPS in vivo. Long-term competitive transplantations with CD201^+^, CD201^−^ or SCA1^+^ LK cells showed that most reconstituting HSCs are within the LK CD201^+^ population after LPS challenge. However long-term competitive repopulation potential of LK SCA1^+^ cells from LPS-treated mice was much more severely reduced than that of LK CD201^+^ cells from the same LPS-treated donors suggesting that the LK SCA1^+^ population in challenged donors becomes contaminated with CD201^−^ progenitors devoid of long-term repopulation potential. Based on CD201 gating strategy, we re-assessed the effect of LPS on HSC and MPP cycling and mobilization, and their dependency on MY88 and TRIF adaptors. In conclusion, CD201 enables a more accurate analysis of mouse HSC and MPP subsets in all inbred strains in septic conditions or steady-state.

**Highlights:**

- SCA1 expression is markedly enhanced on MPPs and myeloid progenitors in vivo following LPS treatment whereas CD201 expression is not.
- Over 90% of the long-term competitive repopulation potential resides within LK CD201+ cells after LPS treatment.
- LK CD201+ cells sorted from LPS-treated mice give 4.4-fold higher multi-lineage chimerism than LK SCA1+ cells.
- SCA1 upregulation leads to overestimation of LT-HSC and MPP subsets, and underestimation of myeloid progenitors in femoral BM of LPS-treated mice.

## INTRODUCTION

The SCA1 (Ly6A/E) antigen has been used for over three decades to identify transplantable mouse hematopoietic stem cells (HSC) and multipotent progenitors (MPP) among lineage-negative (Lin^−^) KIT/CD117^+^ cells [1,2]. However, the SCA1 antigen suffers from 2 major pitfalls. Firstly, although useful to identify HSC in inbred mouse strains of the Ly6A.2 haplotype such as AKR, C57BL, DBA and SJL, SCA1 is poorly expressed by HSC and MPP from Ly6A.1 haplotype strains such as BALB/c, C3H, CB [3] and in Ly6A.2 nonobese diabetic (NOD) strains [4,5], which precludes proper HSC characterization and isolation using SCA1 antibodies in these latter strains. Secondly, SCA1 expression can become strongly induced on more mature lineage-committed hematopoietic progenitor cells (HPC) in inflammatory conditions involving type 1 [6,7] and type 2 [8,9] interferon (IFN) signaling, pathogen-associated molecular patterns (PAMPs) that trigger interferon-mediated responses such as polyinosinic : polycytidylic acid (poly(I:C)) [6,7,10] or lipopolysaccharide (LPS) [11–15], or following bacterial infection and sepsis [6,12,14]. Published analyses of HSC based SCA1 positivity suggest that phenotypic HSC characterized as Lin^−^KIT^+^SCA1^+^CD150^+^ or CD48^−^CD150^+^ or CD34^−^CD150^+^ or side population cells become more proliferative, expand in the bone marrow (BM) and contain more genomic DNA double strand breaks in response to administration of IFN-α [6,7], IFN-γ [8] or PAMPs eliciting an IFN response such as poly(I:C) [6,7] or LPS [12,14–16]. However, interpretation of the results based on conventional phenotypic HSC based on SCA1 positivity following administration of IFNs, PAMPs or sepsis remains uncertain because the HSC/MPP marker SCA1 also becomes highly and promiscuously expressed on many myeloid progenitors which remain otherwise SCA1^−^ in steady-state [10,11]. As a consequence, the Lin^−^KIT^+^SCA1^+^ (LKS^+^) population no longer enriches for HSC and MPP or reflect their biological function in these inflammatory conditions [10,11]. It is therefore critical to find new mouse HSC surface markers whose expression remains stable upon treatment with IFNs or PAMPs to refine these analyses.

For these reasons, we aimed to identify a cell surface marker that more faithfully reflects HSC function even during inflammation or infection across various mouse strains and is also expressed on primate and human HSC to enable interspecies comparisons. We focused on the endothelial protein C receptor CD201, a cell surface marker of reconstituting HSC in all mouse strains [5,17,18] as well as reconstituting human fetal liver and cord blood HSC [19,20]. We now report that unlike SCA1, CD201 expression does not dramatically change on HSC and MPP from C57BL/6 mice following in vivo LPS treatment, enabling a more accurate assessment of the actual effects of LPS-mediated inflammation on proliferation, expansion, and engraftment potential of mouse HSC following inflammatory challenges.

## METHODS

### Mice

C57BL/6, B6.SJL CD45.1^+^, and BALB/c mice were purchased from Animal Resource Centre (Western Australia, Australia) or Ozgene (Western Australia), B6.SJL mice from Ozgene or Australian Bio Resources (New South Wales, Australia). *Myd88*^tm1Aki^ (*Myd88*^−/−^) [21] mice were donated by Dr Antje Blumenthal (The University of Queensland Frazer Institute) and *Ticam1*^tm1Aki^ (*Ticam1*^-/-^) [22] mice by Prof Mark Smyth (QIMR Berghofer Medical Research Institute). All strains had been backcrossed over 10 generations into the C57BL/6 background. Mice were 8-10 week-old. Male mice were used for all the experiments except female mice were used as B6.SJL recipients for long-term transplantation. All experiments were approved by The University of Queensland Animal Experimentation Ethics Committee.

### Treatments

Gamma-irradiated purified LPS from Escherichia coli strain 0111:B4 (Sigma Aldrich catalog # L4391) was administered at 2.5 mg/kg body weight intraperitoneally daily for two consecutive days; control mice were administered with an equivalent volume of saline. Mice were left to rest 48 hours after the last injection before tissue sampling.

### Tissue harvest

At experimental endpoints, mice were anesthetized with isoflurane and 0.5-1mL of blood collected into heparinized EDTA tubes by cardiac puncture before cervical dislocation. The BM of one femur was flushed into 1 mL ice-cold phosphate buffered saline (PBS) containing 2% newborn calf serum (NCS) and 4 mM EDTA using a 1mL syringe mounted with a 21G needle, and BM cells washed in the same buffer during subsequent antibody staining. For BM fluid collection, the BM of one femur was flushed using a syringe containing 1 mL PBS and the cell suspension was centrifuged twice at 370g for 5 mins at 4°C and supernatants collected, aliquoted and stored at -80°C as previously described [23–25].

Before processing hematology indices of blood and BM cells were measured using Mindray BC-5000 Vet Auto hematology analyser (Biomedical Electronics Co. LTD., China).

Blood samples for flow cytometry were first cleaned by erythrocyte lysis by a 5-minute incubation in 5 volumes of 10 mM NaHCO_3_, 150 mM NH_4_Cl, 1 mM EDTA pH=7.4 buffer at room temperature. Blood leukocytes were then washed twice by centrifugation in PBS with 2% NCS.

For plasma collection, blood samples were centrifuged twice at 800g for 10 mins and plasma collected, aliquoted and stored at -80°C.

### Colony assays

Ten µl whole blood aliquots were were seeded in 35 mm petri dishes and covered with 1mL Iscove’s modified Dulbecco’s medium (IMDM) supplemented with 1.6% methylcellulose, 20% FCS and optimal concentrations of conditioned media containing recombinant mouse IL-3, IL-6 and KIT ligand as previously described [26]. Colonies were scored after 10 days of culture at 37 °C in a 5% CO_2_ humidified incubator.

### Bone marrow transplantation

Competitive repopulation assays were used to quantify the long-term repopulating potential of Lin^−^KIT^+^(LK)201^+^, LK201^−^, LKS^+^, LK201^+^FLT3^−^CD48^−^CD150^+^ and LKS^+^FLT3^−^CD48^−^ CD150^+^ cells sorted from the BM of saline- or LPS-challenged donor mice (CD45.2^+^). Eight-week-old C57BL/6 (CD45.2^+^) mice were challenged once daily with saline or 2.5 mg/kg LPS for 2 consecutive days and then left to rest for 48 hours. Donor mice were then euthanized by cervical dislocation, femurs, tibias, and pelvic bones were harvested, cleaned of muscles and crushed in a mortar/pestle containing ice-cold PBS with 2% FCS and 5 mM EDTA. Cells were enriched for KIT/CD117 expressing cells by magnetic-activated cell sorting (MACS) using mouse CD117 microbeads (Miltenyi Biotec) following manufacturer’s instructions. Between 1-2 million KIT^+^-enriched cells from each donor mouse were stained with fluorescein isothiocyanate (FITC)-conjugated lineage (Lin) antibody cocktail (CD3ε, CD5, B220, CD11b, Gr-1 and Ter119), anti–SCA1-PECy7, biotinylated CD201, anti–KIT-allophycocyanin-cyanin7 (APCCy7), CD48-brilliant violet (BV421), CD150-phycoerythrin (PE), anti-FLT3-PECF594 and anti-CD45-BV785. Cells were then washed twice in PBS + 2% NCS + 5mM EDTA and then stained with streptavidin-APC and rewashed. LK201^+^, LK201^−^ or LKS^+^ cells were sorted in PBS containing 20 % FCS using BD FACSAria Fusion sorter (BD Bioscience, San Jose, CA) and profiles for expression of FLT3, CD48 and CD150 were visualized to assess the quality of sorted cells. Equal numbers of sorted donor cells from each mouse of donor experimental group (saline- or LPS-treated) were pooled to form the grafts for the 2 groups. Eight-week-old recipients (B6.SJL CD45.1^+^) were irradiated with two split doses of 5.0 Gy 24 hours before transplantation. Sorted 2,000 LK201^+^, LK201^−^ or LKS^+^ cells were mixed with 4x10^5^ BM cells from naïve UBC-GFP (CD45.2^+^) mice and transplanted into B6.SJL recipients. For transplants with sorted LK201^+^ FLT3^−^CD48^−^CD150^+^ and LKS^+^ FLT3^−^CD48^−^CD150^+^ HSCs from LPS-treated C57BL/6 donors, 1,340 sorted cells were transplanted per recipient in competition with 4x10^5^ BM cells from naïve B6.SJL (CD45.1^+^) mice into lethally irradiated B6.SJL recipients.

Multi-lineage chimerism was measured by flow cytometry at 4-weeks intervals post-transplant following staining of blood leukocytes with CD3ε-BV421, CD45.1-PE, CD45.2-APC, CD11b-peridinin-chlorophyll-protein-CY5.5 (PercpCY5.5) or CD11b-BV510, anti-B220-APCCY7, LY6G-PECY7 and for GFP for competing cells. Fixable viability stain (FVS)700 (BD Bioscience) was added to all stained samples for dead cell exclusion.

Repopulating units in each recipient mouse were calculated at 20 weeks post-transplantation using the following formula as previously described [27,28] with 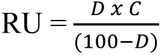 where D is the percentage of donor CD45.2^+^ GFP^−^B and myeloid cells, and C is the number of competing CD45.1^+^ BM repopulation units co-transplanted with the donor cells (C=4 for 400,000 competing BM cells) [27]. Repopulation units were calculated per 2,000 transplanted donor LK201^+^, LK201^−^ or LKS^+^ cells or per 1,340 LK201^+^ FLT3^−^CD48^−^CD150^+^or LKS^+^ FLT3^−^CD48^−^CD150^+^ HSCs.

### Flow cytometry analyses

All fluorescent antibodies used in this study are listed in Supplemental Table 1. To quantify SCA1 expression in the BM and number of HSPC in the BM and blood, 5x10^6^ BM and blood leukocytes were stained in suspension on ice for 40 minutes in mouse CD16/CD32 hybridoma 2.4G2 supernatant containing PercpCY5.5-conjugated lineage (Lin) antibody cocktail (CD3ε, CD5, B220, CD11b, Gr-1, Ter119), anti-SCA1-FITC, anti-KIT-APCCY7, biotinylated CD201, CD48-BV421, CD150-PE, anti-FLT3-PECF59 and CD45-BV785. Cells were then washed twice in PBS + 2% NCS + 5mM EDTA and then stained with streptavidin-APC and rewashed. Fixable viability stain (FVS)700 (BD Bioscience) was added to all stained samples for dead cell exclusion.

To quantify number of BM myeloid progenitors, 5x10^6^ BM cells were stained in MACS buffer (Miltenyi Biotec) containing FITC-conjugated Lin antibody cocktail (CD3ε, CD5, B220, CD11b, Gr-1, Ter119), anti-SCA1-BV510, anti-KIT-APCCY7, biotinylated CD201, CD41-APCCY7, CD16/32-PercpCY5.5, CD105-PE, CD48-BV421, CD150-PECY7, anti-FLT3-PECF59 and CD45-BV785. Cells were then washed twice in PBS + 2% NCS + 5mM EDTA and then stained with streptavidin-APC and rewashed. FVS700 (BD Bioscience) was added to all stained samples for dead cell exclusion. Samples were analyzed either on a Cytoflex (Beckman Coulter) flow cytometer equipped with 640nm, 561nm, 488nm, and 405nm lasers. Uncompensated FCS files were analyzed using FlowJo10 software following compensation with single color antibody stains (Tree Star, Ashland, OR) following post-hoc compensation with single color controls.

### Cell cycle analyses

Femurs, tibias, and pelvic bones were harvested from saline or LPS treated wild type mice and crushed in a mortar/pestle containing ice-cold PBS with 2% NCS and 5mM EDTA and stained for cell-surface antigens and then for Ki67 antigen and DNA content as described previously [29]. Briefly, 1 million KIT^+^-enriched cells were stained with PercpCY5.5-conjugated lineage antibodies (CD3ε, CD5, B220, CD11b, Gr-1 and Ter119), anti–SCA1-PECy7, biotinylated CD201, anti–KIT-APCCY7, CD48-AlexaFluor700, anti-FLT3-PECF594 and CD150-PE. Labeled cells were washed twice in PBS + 2% NCS + 5mM EDTA, then stained with streptavidin-APC and rewashed with PBS, and fixed and permeabilized using the Fix perm cell permeabilization kit (Life Technologies). FITC-conjugated mouse anti–Ki67 was added to the permeabilization buffer and cells were stained for 30 minutes. After washing, cells were incubated in 1 mL of PBS containing 1 µg/mL of RNase A, 0.05% saponin, and 25 µM Hoechst33342 for 10 minutes before analysis. Data were acquired on a Fortessa X20 flow cytometer equipped with 355 nm, 405 nm, 488 nm, 561 nm and 637 nm excitation lasers.

### ELISA

Mouse CXCL12 was quantified using specific ELISA kits (R&D Systems; cat # MCX120) in BM fluids following manufacturer’s instructions. Plates were read with plate reader (Thermo Fisher Scientific) according to manufacturer’s instructions.

### Cytokine measurements with Legendplex bead arrays

Cytokine concentrations were measured in diluted plasma (1:2) using LEGENDplex^TM^ mouse inflammation panel (13 plex; BioLegend, catalog # 740446) following manufacturer’s instructions. Cytokine data were acquired on a CytoFLEX flow cytometer. Cytokine concentrations were analysed using BioLegend’s LEGENDplex™ data analysis software (BioLegend).

### Quantitative RT-PCR

RNA was extracted using GeneJet RNA Clean up and Concentration Micro Kit as specified by manufacturer (ThermoFisher Scientific). cDNA was synthesized with SensiFast cDNA synthesis kit (Alexandria NSW, Australia). qRT-PCR was performed with TaqMan Fast Advanced Mix (ThermoFisher Scientific) with mouse *Hprt* (Mm03024075_m1), *Vcam1* (Mm01320970_m1) primer / probe sets (ThermoFisher Scientific) using QuantStudio^TM^ Real-Time PCR system (ThermoFisher Scientific).

### Statistical analyses

Differences between treatment groups were calculated using GraphPad Prism v8.0.1 software using non-parametric Mann-Whitney test for pairwise comparisons or one-way or two-way ANOVA for multiple comparisons as specified in figure legends.

## RESULTS

### LPS strongly upregulates SCA1 antigen on mouse HSPC in vivo but with little effect on CD201

To evaluate the effect of sepsis on SCA1 and CD201 expression, mice were injected with LPS at 2.5 mg/kg daily or saline for two consecutive days (Figure 1A). LPS significantly increased inflammatory cytokines in blood including type 1/2 IFNs (Supplemental Figure 1) which have been reported to upregulate SCA1 expression on BM Lin^−^ cells [6–9].

**Figure 1.**
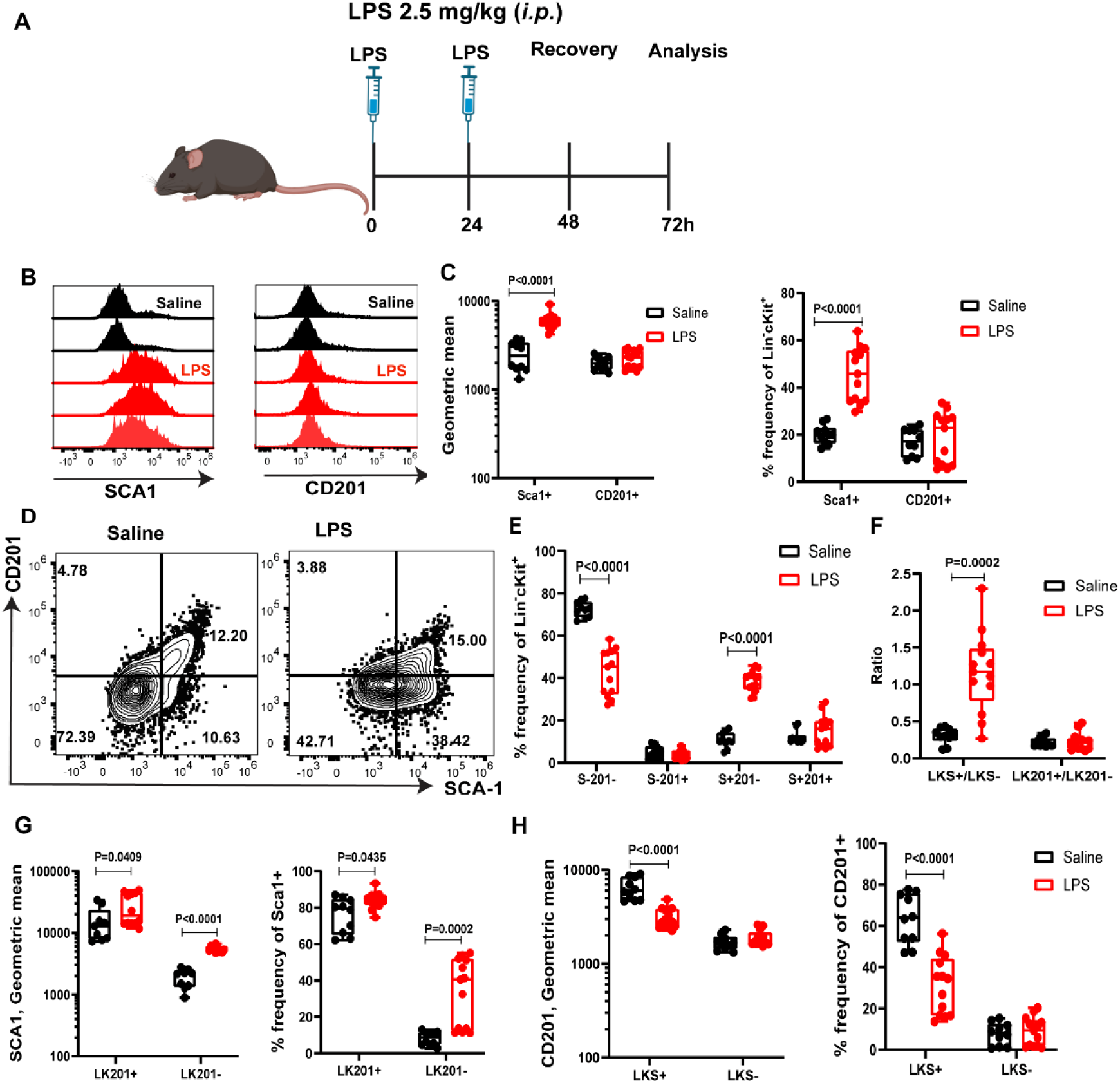
LPS upregulates SCA1 but not CD201 expression on mouse bone marrow HSPC. (**A**) C57BL/6 mice were injected once daily for 2 consecutive days with saline or LPS and tissues harvested after 2 days of recovery. (**B**) Flow cytometry histogram plots showing increased SCA1 but invariant CD201 expression on BM Lin^−^ KIT^+^ cells upon LPS (2.5 mg/kg) or saline treatment. (**C**) Geometric mean and frequency of SCA1^+^ and CD201^+^ cells gated on BM Lin^−^KIT^+^ cells from mice treated with LPS or saline. (**D**) Representative flow cytometry contour plots showing CD201 and SCA1 expression on BM Lin^−^ KIT^+^ cells following LPS or saline treatment. (**E**) Frequencies of SCA1^+^CD201^−^ (S^+^201^−^), SCA1^−^CD201^+^ (S^−^201^+^) and SCA1^+^CD201^+^ (S^+^201^+^) cells on BM Lin^−^KIT^+^ cells after LPS or saline treatment. (**F**) Ratio of LKS^+^/LKS^−^ and LK201^+^/LK201^−^ cells after LPS or saline treatment. (**G**) Fluorescence geometric mean intensity and frequencies of SCA1 expression on LK201^+^ and LK201^−^ cells. (**H**) Fluorescence geometric mean intensity and frequencies of CD201^+^ cells within LKS^+^ and LKS^−^ populations. Data are from three pooled independent experiments performed within a 1-year interval. Each dot is a separate mouse with mean ± SD. Statistical significance was determined using Mann-Whitney test between saline and LPS treated mice.

LPS treatment dramatically increased SCA1 geometric mean fluorescence intensity and frequency of SCA1-positive cells by ∼3-fold and ∼2.5-fold respectively on LK cells (Figure 1B-C) consistent with previous reports [6–9]. In sharp contrast, CD201 expression on LK cells remained unchanged upon LPS treatment with no significant change in CD201 fluorescence geometric mean and no significant increase in the proportion of CD201^+^ cells (Figure 1B,C).

In plots of SCA1 vs CD201 expression on LK cells, LPS dramatically increased 4.3-fold the proportion of CD201^−^SCA1^+^ cells (Figure 1D-E). This increase in LK CD201^−^SCA1^+^ cell proportion was compensated by equivalent decrease in LK CD201^−^SCA1^−^cells whereas the proportion of LK CD201^+^SCA1^+^ cells, which has been reported to contain apex tip HSC [30], was not significantly altered by LPS treatment (Figure 1D-E). As a consequence of this shift in SCA1 expression, LPS treatment dramatically enhanced the ratio of LKS^+^/LKS^−^ cells in the BM whereas the ratio of LK201^+^/LK201^−^ remained unchanged (Figure 1F). Additionally, we measured how LPS treatment altered SCA1 expression on LK201^+^ and LK201^−^ cells (Figure 1G) and conversely how it altered CD201 expression on LKS^+^ and LKS^−^ cells (Figure 1H). LPS significantly increased SCA1 expression in both LK201^+^ and LK201^−^ cells whereas LPS significantly decreased CD201 average expression on LKS^+^ cells only. Altogether, these data suggest that LPS challenge increased SCA1 expression on LK cells but not CD201 expression, resulting in a dilutive effect on the average CD201 expression within the LKS^+^ population in LPS-treated mice. This is consistent with a recent study showing that most LKS^−^ cells become SCA1^+^ following systemic infection [11]. Overall these data show that CD201 expression is more stable on LK cells upon LPS challenge.

We next analyzed SCA1 and CD201 expression on phenotypic HSC and MPP subsets that were gated omitting SCA1 and CD201 for the gating, and then examined how these two markers evolve in these subsets in response to LPS (Figure 2). LT-HSC were gated as LK FLT3^−^CD48^−^CD150^+^, ST-HSC as LK FLT3^−^CD48^−^CD150^−^, MPP2 as LK FLT3^−^CD48^+^CD150^+^, MPP3 as LK FLT3^−^CD48^+^CD150^−^ and MPP4 as LK FLT3^+^CD48^+^CD150^−^ (Supplemental Figure 2). LPS treatment in vivo dramatically increased SCA1 expression on LT-HSC and MMP2-4 but not on ST-HSC (Figure 2A-J). In contrast, LPS did not increase CD201 expression on LT-HSC, ST-HSC, and MPP4 populations but induced only a small but significant increase in CD201 mean fluorescence intensity on MPP2 and MPP3 (Figure 2A-J). The consequence of this SCA1 upregulation on MPPs and more committed myeloid progenitors (see paragraph below), was that the number of “apparent” SCA1^+^ MMP2 and MPP3 per femur appeared to dramatically increase in response to LPS whereas the changes in LK201^+^ MPP2 and MPP3 per femur were more moderate (Figure 2K,L and Supplemental Table 2). Likewise, the marked decrease in LK201^+^ MPP4 numbers per femur in response to LPS was not as marked as that of LKS^+^ MPP4. All these effects on the number of CD201^+^ versus SCA1^+^ MPPs per femur in response to LPS can be explained by the marked SCA1 up-regulation on these MPP subsets (Figure 2H-J).

**Figure 2.**
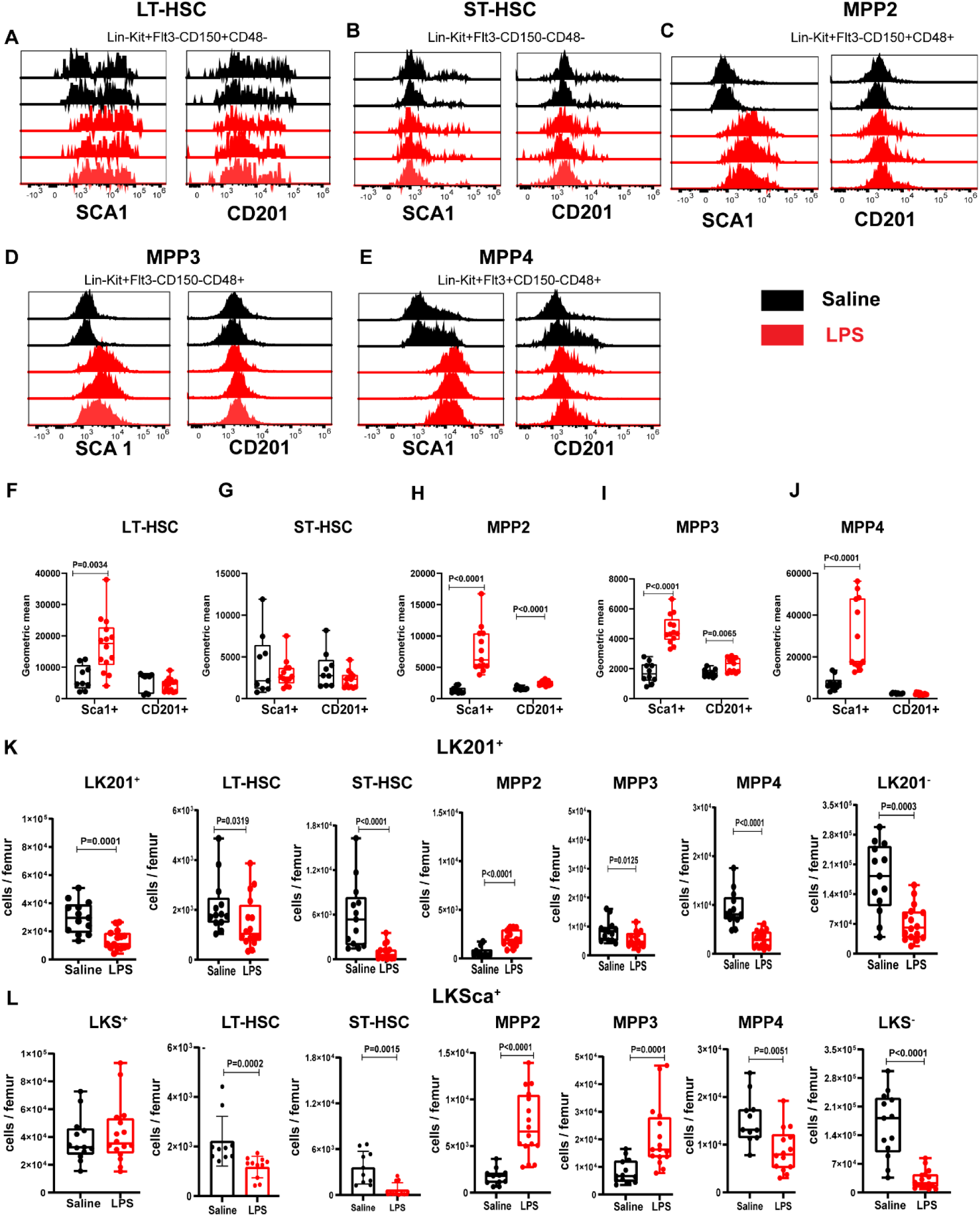
LPS upregulates SCA1 expression on bone marrow phenotypic HSC and MPP populations. C57BL/6 mice were injected twice with LPS once daily and then rested for 48 hours as in Figure 1A. (**A-E**) Overlays of SCA1 and CD201 fluorescence histograms in (**A**) LK FLT3^−^CD150^+^CD48^−^ long term reconstituting (LT) HSCs, (**B**) LK FLT3^−^CD150^−^CD48^−^ short term reconstituting (ST) HSCs, (**C**) LK FLT3^−^CD150^+^CD48^+^ MPP2, (**D**) LK FLT3^−^CD150^−^CD48^+^ MPP3 and (**E**) LK FLT3^+^CD150^−^CD48^+^ MPP4. The black profiles are from 2 representative saline-treated mice, the red profiles from 3 representative LPS-treated mice. (**F-G**) Geometric fluorescence mean intensities of SCA1 and CD201 on (**F**) LT-HSC, (**G**) ST-HSC, (**H**) MPP2, (**I**) MPP3 and (**J**) MPP4 as defined above following treatment with saline (black) or LPS (red). (**K**) Number of cells per femur in HSC and MPP subsets in response to LPS using CD201 to define these subsets. (**L**) Number of cells per femur in HSC and MPP subsets in response to LPS using SCA1 to define these subsets. Each dot is a separate mouse. Statistical significance was calculated using Mann-Whitney test.

In respect to downstream myeloid progenitors, LPS has also been reported to dramatically reduce the numbers of common myeloid progenitors (CMP) and granulocyte-macrophage progenitors (GMP) in the BM [14]. These measurements were based on SCA1^−^gating according to their phenotype in steady-state BM [14]. Therefore, we assessed SCA1 expression on CMP, GMP, myeloid-erythroid progenitor (MEP), megakaryocyte progenitors (MegP), burst-forming units erythroid (BFU)-E and colony-forming units (CFU)-E based on the LK201^−^ population (gating strategy in Supplemental Figure 3) as CD201 expression is stable following LPS challenge (Figure 1C,F,H). LPS significantly upregulated SCA1 expression on CMP, GMP, MEP, MegP and to a lower extent BFU-E with increased mean fluorescence intensity. Similarly, LPS challenge caused a larger proportion of LK201^−^progenitor cells to become SCA1^+^ (Supplemental Figure 4A-C). As a consequence, the apparent reduction in the number of CMP, MEP, MegP and BFU-E per femur in response to LPS measured when LKS^−^ gate was used as in previous reports [14] was not as pronounced when these progenitors were gated from LK201^−^ cells instead of LKS^−^ cells (Supplemental Figure 4D,E and Supplemental Table 3).

These results confirm that upon LPS treatment, SCA1 expression is increased on LT-HSC, MPP2-4, as well as myeloid and megakaryocyte progenitors while CD201 expression remains more stably expressed following LPS exposure. Therefore, CD201 appears to be a more appropriate marker than SCA1 to delineate HSPC subsets following sepsis, LPS and presumably IFN-mediated challenges.

To determine whether these effects of LPS were restricted to C57BL/6 strain, SCA1 and CD201 expression on HSPC from BALB/c mice, a Ly6A.1 haplotype strain well known to poorly express SCA1 antigen [3], was also measured. Following the same gating strategy as in Figure 1, we found that LPS strongly up-regulated SCA1 expression whereas CD201 expression was less affected in BALB/c mice (Supplemental Figure 5A-F). When HSC and MPP were gated from the LK201^+^ subset, SCA1 expression was strongly up-regulated by LPS treatment on LT-HSC and MPP2 but only mildly on ST-HSC, MPP3 and MPP4 (Supplemental figure 5G-K). Thus, SCA1 upregulation in response to LPS is not restricted to strains of the Ly6A.2 haplotype such as C57BL/6. The small increase in CD201 expression observed in LPS-treated mice may reflect an actual expansion of HSC and MPP compartments in these mice.

### Long-term reconstituting HSCs are more frequent in the LK201^+^ population than in the LKS^+^ population in septic conditions

Our data show that LPS treatment strongly induces SCA1 expression on myeloid progenitors, cells that under normal steady-state conditions are SCA1^−^. This observation further suggests that the reported reduction in hematopoietic reconstitution potential of LKS^+^ cells upon LPS treatment [11,14] could be due in part to contamination of this population with temporarily SCA1 expressing myeloid progenitors which do not engraft long-term. To test this hypothesis, a side-by-side comparison of LK201^+^ versus LKS^+^ cell engraftment potential following strong inflammatory challenge such as LPS was performed. C57BL/6 CD45.2^+^ donor mice were first challenged with LPS and then 2,000 sorted BM LK201^+^, LK201^−^ or LKS^+^ were transplanted into lethally irradiated congenic CD45.1^+^ recipients in competition with 4x10^5^ whole BM GFP^+^ cells (from *Ubc*-GFP CD45.2^+^) mice. Following LPS challenge, sorted donor LK201^+^ cells exhibited superior myeloid, B and T lineage reconstitution (Figure 3 and gating strategy in Supplemental Figure 6) with a 4.4-fold higher reconstitution potential compared to LKS^+^ cells sorted from the same donors as determined by repopulation unit (RU) content (Figure 3F). Overall, post-transplant reconstitution of sorted donor LK201^−^ cells was poor approximately 75-fold lower than LK201^+^ cells (Figure 3F). Together, the LK201^+^ population remains enriched in competitive long-term repopulating HSC compared to LKS^+^ cells due to reduced contamination from lineage-committed progenitors that have become SCA1^+^ following LPS challenge.

**Figure 3.**
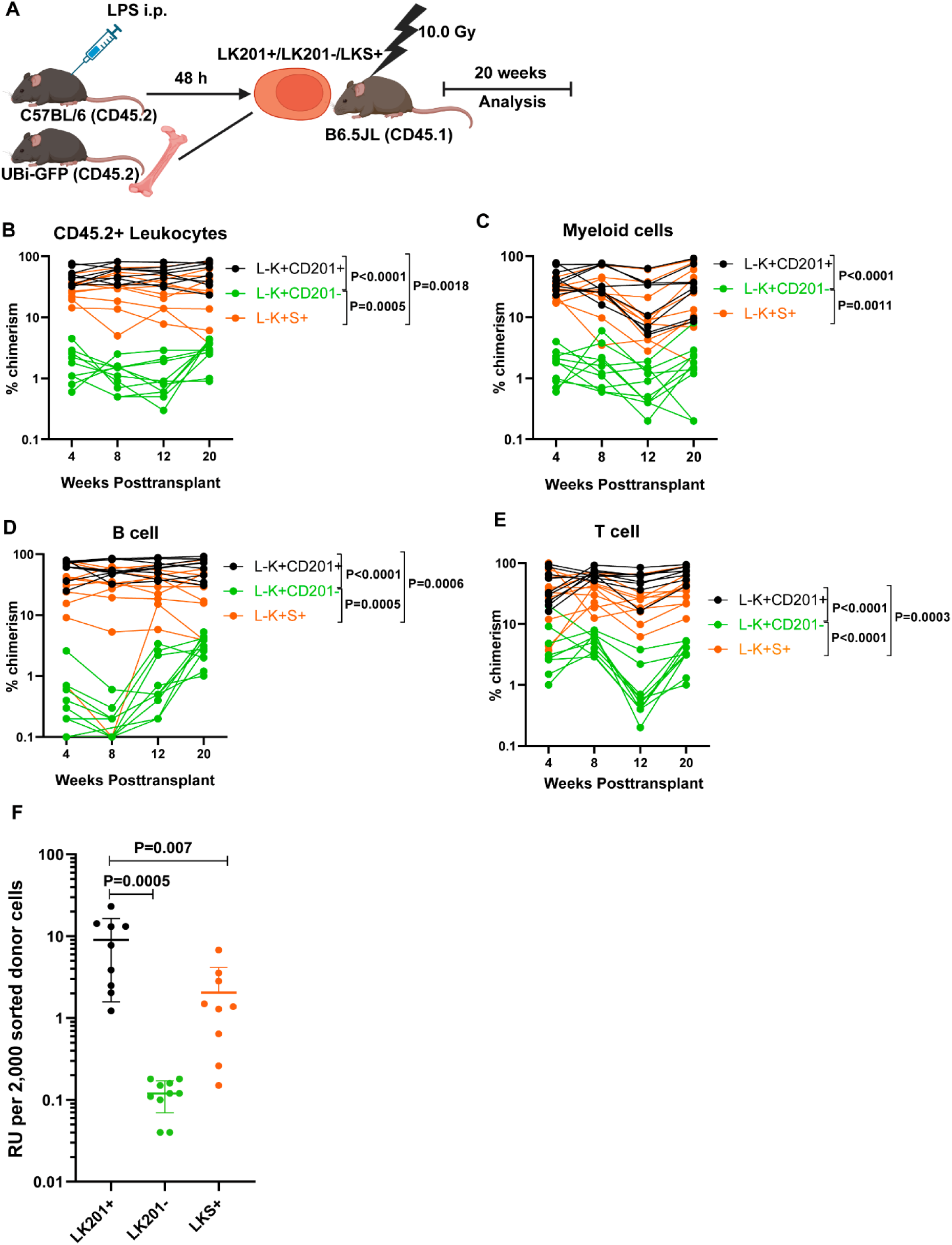
Lin^−^ KIT^+^ CD201^+^ cells contain most long-term competitive repopulating activity after LPS treatment. **(A**) Schematic of long-term competitive repopulation assay. Donor C57BL/6 mice were injected twice with LPS once daily and then rested for 48 hours as in Figure 1A. Two thousand LK201^+^, LK201^−^ and LKS^+^ cells sorted from the BM and transplanted into lethally irradiated B6.SJL CD45.1^+^ recipients in competition with 400,000 BM cells from UBI-GFP mice. Recipient mice were maintained for 20 weeks and blood collected every 4 weeks. The figure was created with BioRender.com. Kinetics of engraftment of (**B**) donor GFP^−^CD45.2^+^ leukocytes, (**C**) donor GFP^−^CD45.2^+^ CD11b^+^ myeloid cells, (**D**) GFP^−^CD45.2^+^ B220^+^ B cells and (**E**) GFP^−^CD45.2^+^ CD3^+^ T cells in the blood as measured by flow cytometry every 4 weeks post-transplantation. Dots corresponding to each individual recipient are connected by a continuous line. P values were calculated using two-way ANOVA with Sidak’s multiple comparison test. (**F**) Repopulation unit (RU) were calculated for 2,000 donor cells upon 20 weeks of transplantation. Each dot is a separate mouse with mean ± SD. Statistical significance was calculated using one-way ANOVA with Sidak’s multiple comparison test.

A recent study suggests that both CD201^+^ and CD201^−^ populations contain HSC activity during inflammation with different engraftment output [13]. We compared the competitive repopulation potential of 2,000 sorted LK201^+^ and LK201^−^ cells from donor mice in steady-state or following LPS challenge (Figure 4), donor LK201^−^ HSCs led to very poor chimerism in all lineages with a 93- to 115-fold lower RU content compared to LK201^+^ cells. In contrast, despite containing the highest repopulation potential, LK201^+^ from LPS-treated donors showed reduced chimerism in all lineages compared to LK201^+^ cells from saline-treated donors with RU content decreasing 4.0-fold (Figure 4E). Together, these results indicate that LPS treatment reduces engraftment potential of LK201^+^ cells, however LK201^+^ cell repopulation potential remained 4.4-fold higher than that of LPS-challenged LKS^+^ cells (Figure 3F). Importantly, the reconstituting HSC pool remains almost wholly contained within cells of the LK201^+^ phenotype.

**Figure 4.**
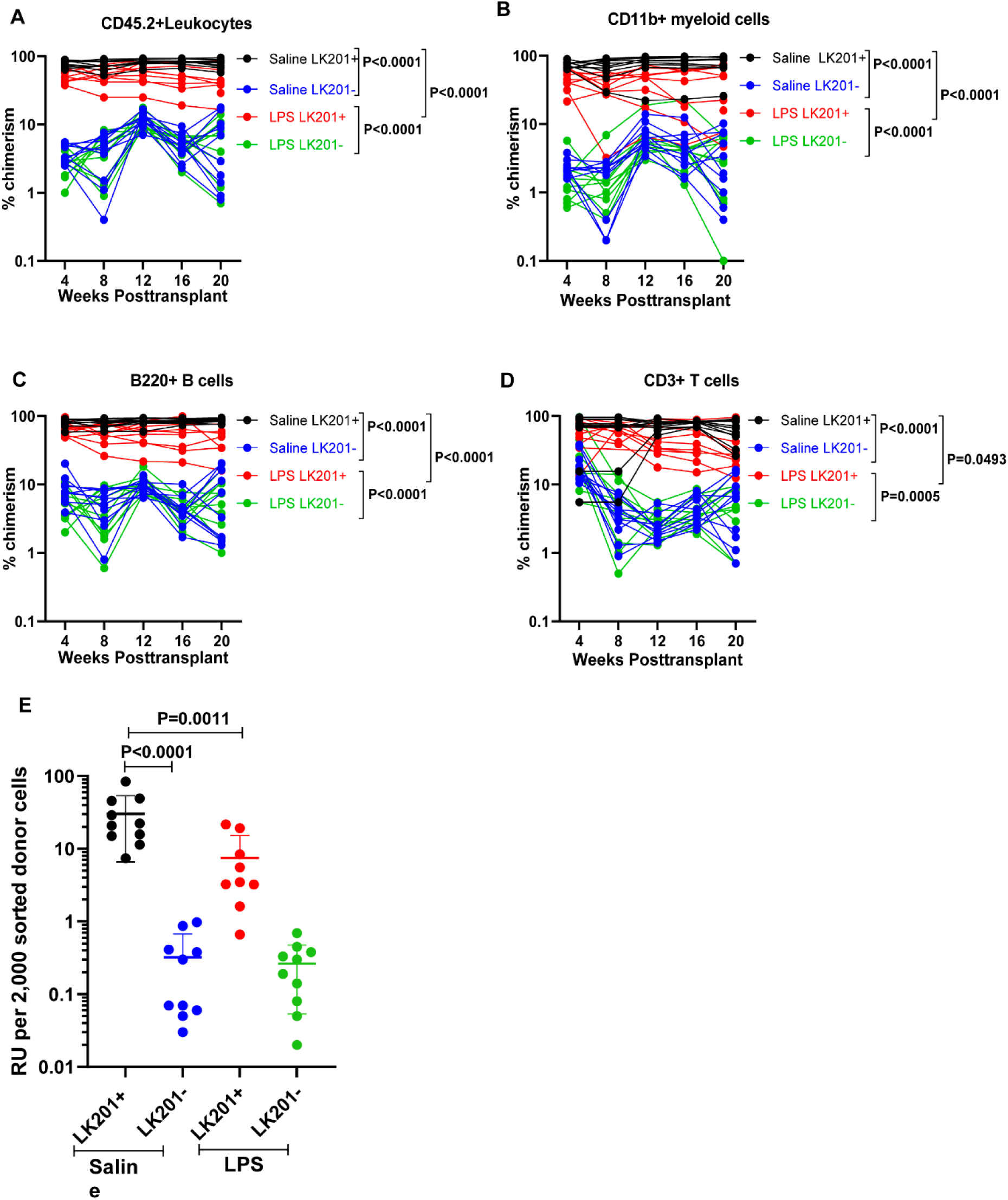
Lin^−^ KIT^+^ CD201^+^ HSC contain most long-term competitive repopulating activity in steady-state or after LPS challenge. Donor C57BL/6 mice were injected once daily with saline or LPS for 2 consecutive days and harvested after two days of recovery as in figure 1A. Two thousand BM LK201^+^ and LK201^−^ cells were sorted and transplanted into lethally irradiated B6.SJL CD45.1^+^ mice in competition with 400,000 BM cells from UBI-GFP mice and maintained for 20 weeks with blood collection every 4 weeks via tail vein. Kinetics of engraftment of (**A**) donor GFP^−^CD45.2^+^ leukocytes, (**B**) donor GFP^−^CD45.2^+^ CD11b^+^ myeloid cells, (**C**) GFP^−^CD45.2^+^ B220^+^ B cells and (**D**) GFP^−^CD45.2^+^ CD3^+^ T cells in the blood as measured by flow cytometry every 4 weeks post-transplantation. In panel A-E, Dots corresponding to each individual recipient are connected by a continuous line. P values were calculated using two-way ANOVA with Sidak’s multiple comparison test. (**E**) Repopulation unit (RU) were calculated for 2,000 donor cells upon 20 weeks of transplantation. Each dot represents a separate recipient mouse. Bars are mean ± SD. Statistical significance was calculated using one-way ANOVA with Sidak’s multiple comparison test. Black: Recipients of LK201^+^ cells from saline-treated donors; Red: recipients of LK201^+^ cells from LPS-treated donors; Blue: recipients of LK201^−^ cells from saline-treated donors; Green: recipients of LK201^−^ cells from LPS-treated donors.

To determine whether the increased loss of long-term engraftment potential of conventional LKS^+^ cells after in vivo LPS was due to dilution of this population by myeloid progenitors or due to HSC intrinsic mechanisms, we sorted LKS^+^ and LK201^+^ phenotypic LT-HSC (FLT3^−^CD48^−^CD150^+^) from the same LPS-treated donors and performed side-by-side comparison in a competitive repopulation assay. Cells from both phenotypes contained similar multi-linage long-term reconstitution potential (RU) but LK201^+^FLT3^−^CD48^−^CD150^+^ phenotypic LT-HSC had significantly better B cell lineage reconstitution compared to LKS^+^FLT3^−^CD48^−^CD150^+^ phenotypic LT-HSC from LPS-treated mice (Figure 5).

**Figure 5.**
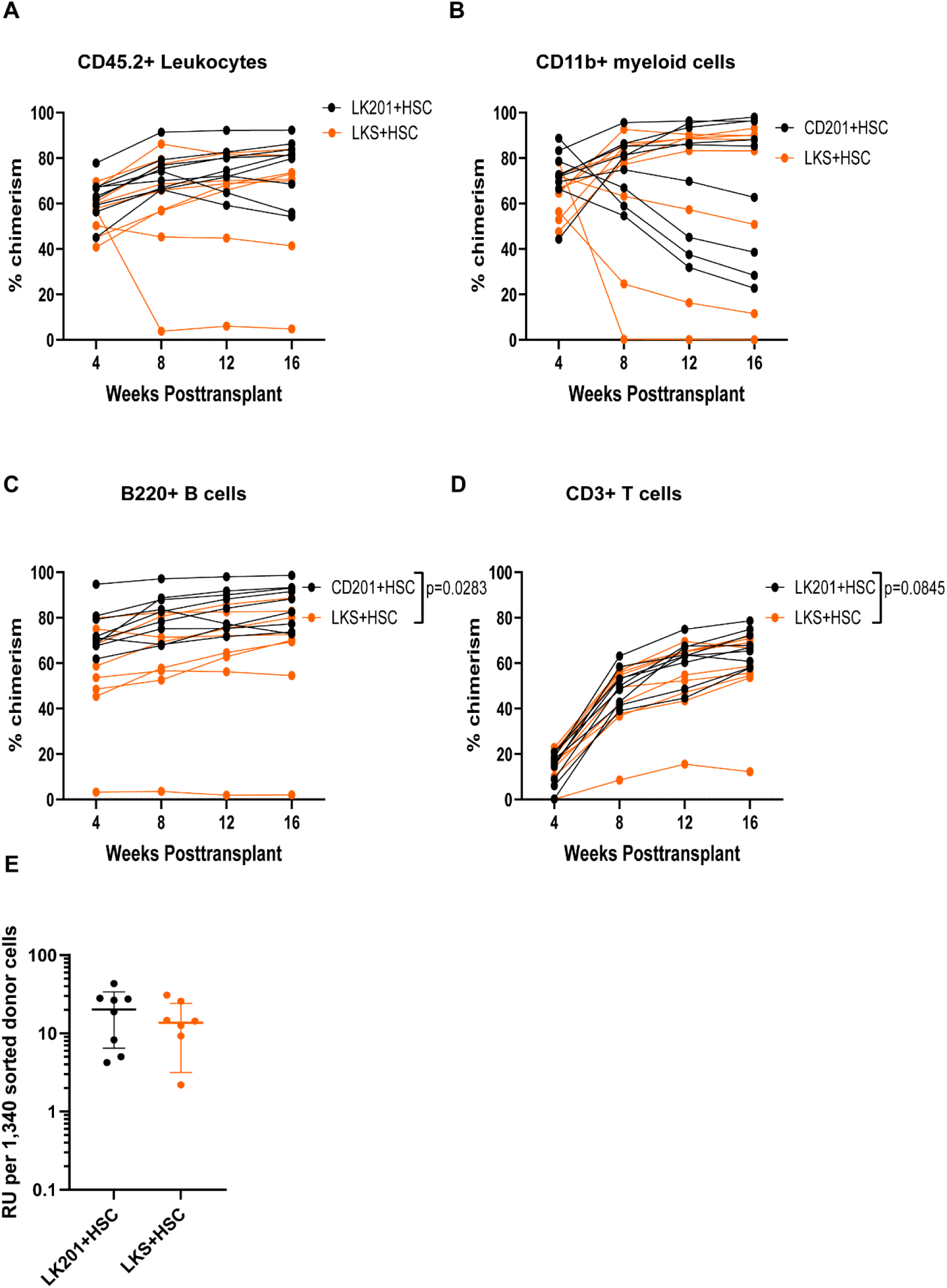
Phenotypic CD201^+^ LT-HSC from LPS-treated mice have superior self-renewal capacity than SCA1^+^ LT-HSC in secondary recipients. All donor C57BL/6 mice were treated with LPS as in Figure 3A. 1,340 sorted LK201^+^ FLT3^−^ CD48^−^ CD150^+^ and LKS^+^ FLT3^−^ CD48^−^CD150^+^ cells were transplanted in competition with 400,000 BM cells from naïve B6.SJL mice into lethally irradiated primary recipients which were bled every 4 weeks to following engraftment. Kinetics of engraftment in primary recipients of (**A**) donor CD45.2^+^ leukocytes, (**B**) donor CD45.2^+^ CD11b^+^ myeloid cells, (**C**) CD45.2^+^ B220^+^ B cells and (**D**) CD45.2^+^ CD3^+^ T cells in the blood as measured by flow cytometry every 4 weeks post-transplantation. P values were calculated using two-way ANOVA with Sidak’s multiple comparison test. **(E**) Repopulation unit (RU) were calculated for 1,340 donor cells 16 weeks post-transplantation. Each dot represents a separate recipient mouse. Bars are mean ± SD. Black: recipients of LK201^+^ LT-HSC. Red: recipients of LKS^+^ LT-HSC.

Collectively, these data support the notion that LPS treatment leads to temporary SCA1 upregulation on myeloid progenitors and an apparent dilution of the LKS^+^ compartment with SCA1^+^ CD201^−^ lineage-committed progenitors which are devoid of long-term repopulation potential. Thus, a gating strategy that includes CD201^+^ cells instead provides better delineation of HSC, MPP and myeloid progenitor subsets in mice challenged with LPS.

### LPS enhances CD201^+^ HSC and MPP proliferation

Approximately 60% phenotypic HSC are reportedly quiescent in G0 phase of the cell cycle in steady-state [31] and exposure to LPS enhances their proliferation [12,14,15]. However in all these studies, SCA1 positivity was used to gate HSC and MPP. CD201^−^ lineage-committed progenitors which actively divide even in steady-state conditions [31,32], we re-assessed the effect of LPS treatment on HSC and MPP cycling based on LK201^+^ phenotype and compared their cycling status to HSC and MPP subsets based on LKS^+^ phenotypes (gating strategy in Supplemental Figure 7).

Using the LKS^+^ cell population gating, the proportion of quiescent LT-HSC, ST-HSC and MPP4 in G0 significantly decreased following LPS administration whereas the proportion in G1 phase significantly increased (Figure 6A-E) consistent with previous reports [12,14,15]. LPS treatment did not alter SCA1^+^ MPP2 and MPP3 cycling status.

**Figure 6.**
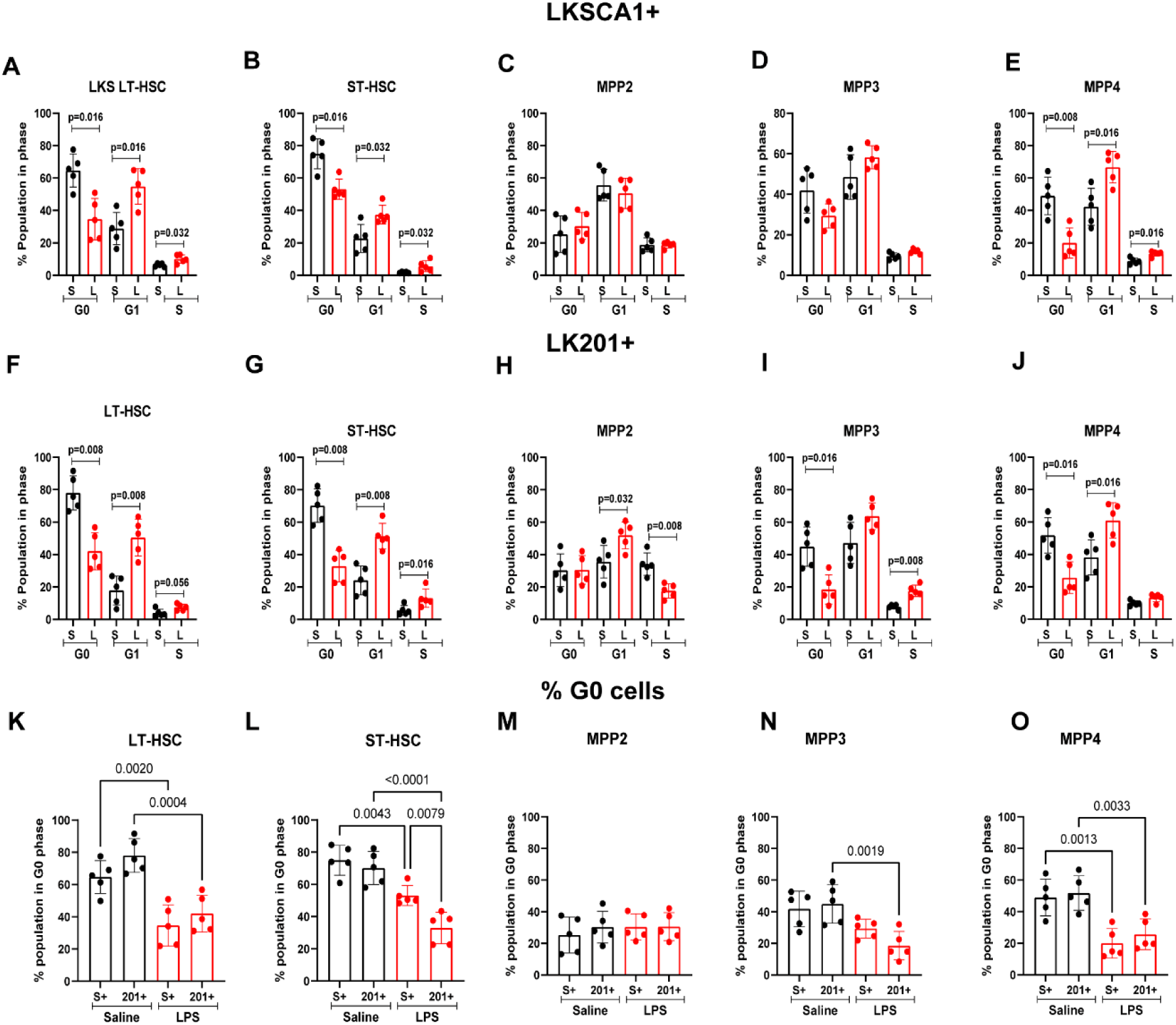
LPS treatment increases proliferation of BM CD201^+^ HSCs in vivo. C57BL/6 mice were injected once daily with saline (black dots columns) or LPS (red dots columns) for 2 consecutive days and harvested after two days of recovery. (**A-E**) Top row showing proportion of phenotypic LKS^+^FLT3^−^CD150^+^CD48^−^ long term reconstituting (LT) HSCs, LKS^+^ FLT3^−^CD150^−^CD48^−^ short term reconstituting (ST) HSCs, LKS^+^ FLT3^−^CD150^+^CD48^+^ MPP2, LKS^+^ FLT3^−^CD150^−^CD48^+^ MPP3 and LKS^+^FLT3^+^CD150^−^CD48^+^ MPP4 in phases G0, G1 and S/G2/M of the cell cycle. (**F-J**) Bottom row shows proportion of LK201^+^FLT3^−^CD150^+^CD48^−^LT-HSCs, LK201^+^FLT3^−^CD150^−^CD48^−^ ST-HSCs, LK201^+^ FLT3^−^CD150^+^CD48^+^ MPP2, LK201^+^ FLT3^−^CD150^−^CD48^+^ MPP3 and LK201^+^FLT3^+^CD150^−^CD48^+^ MPP4 from saline and LPS treated mice in phases G0, G1 or S/G2M of the cycle. (**K-O**) Side by side comparison in the number of quiescent cells in G0 among CD201^+^ or SCA1^+^ phenotypic LT-HC, ST-HSC, MPP2, MPP3 and MPP4 as defined above. Each dot represents a separate mouse. columns are mean ± SD. Statistical significance was determined using Mann-Whitney test between saline and LPS treated mice for panels A-J, and by one-way ANOVA for panels K-O.

When HSPC were defined using LK201^+^ phenotype, LPS administration induced similar significant decrease in HSC, and MPP4 proportion in G0 cell cycle, a drop in quiescent ST-HSC in cycle, and a significant LPS-mediated drop in quiescent MPP3 which not apparent when LKS^+^ gating used.

Taken together these data indicate that based on LKCD201^+^ gating to define HSC and MPP subsets, LPS treatment in vivo increase recruitment into cell cycle of all HSC and MPP subsets except MPP2.

### Expression of both TRIF and MYD88 is required for SCA1 upregulation in response to LPS

LPS binds to Toll-like receptor 4 (TLR4) and activates NF-κB and IFN signaling pathways through two different adaptors: myeloid differentiation factor 88 (MYD88) and TIR-domain containing adapter-including interferon-β (TRIF also called TICAM-1) [21,22]. To determine which of these adaptor molecules were involved in LPS-induced SCA1 upregulation, LPS was injected in mice defective for either TRIF (*Trif^-/-^*) or MYD88 (*Myd88^-/-^*). In either *Trif^-/-^* or *Myd88^-/-^*mice, SCA1 was not up-regulated on LK cells following in vivo challenge with LPS either in frequency of SCA1^+^ cells or SCA1 mean fluorescence intensity (Supplemental Figure 8A-B). This suggests LPS increases SCA1 expression requires both TRIF and MYD88 together.

### Expression of both TRIF and MYD88 is required for HSPC mobilization into blood

Using SCA1 as HSPC marker, we and others previously have shown that LPS promotes HSPC mobilization [33,34]. However to what extent phenotypic HSC mobilization has been overestimated due to SCA1 upregulation on myeloid progenitors is an unanswered question. We found that LPS enhanced mobilization of CFC, LK201^−^, LKS^−^, LK201^+^, LKS^+^ into the blood (Supplemental Figure 9) in line with the increase in blood WBC counts (Supplemental Figure 10) although LK201^+^ cell mobilization was 3.6-fold lower than LKS^+^ cell mobilization (p=0.0003 Mann-Whitney test, Supplemental Figure 9D,E). Despite this, mobilization of phenotypic LT-HSC based on either CD201 or SCA1 positivity were similar (Supplemental Figure 9F,G). This suggests that when phenotypes that involve SCA1 positivity are used, they lead to overestimation of MPP mobilization in response to LPS.

LPS did not mobilize CFC or HSPC into the blood in mice defective for either TRIF or MYD88 showing that signaling through both adaptors is required for mobilization in response to LPS (Supplemental Figure 9). In line with this, LPS increased WBC, monocytes, neutrophil, eosinophil, and basophils in the blood in a manner that also required both MYD88 and TRIF (Supplemental Figure 10). LPS also significantly reduced the number of lymphocytes and platelets via both TRIF and MYD88 (Supplemental Figure 10).

We finally measured the effect of LPS on the expression of CXCL12 chemokine and cell adhesion molecule VCAM1, which are expressed by niche cells and retain HSPC within the BM [23–25,35,36]. Unexpectedly, the suppressive effect of LPS on CXCL12 protein and *Vcam1* transcript expression in the BM was only dependent on MYD88 with no effect of TRIF deletion (Supplemental Figure 9H,I).

## DISCUSSION

In a model of sepsis, we report that unlike SCA1, CD201 expression is not dramatically upregulated on HSPC in response to LPS treatment in vivo (Figures 1-2, Supplemental Figure 4). We found that most long-term reconstitution activity was contained within the LK201^+^ population in steady-state or following LPS treatment and that the repopulating activity following LPS treatment remains 4-fold higher in sorted LK201^+^ cells than LKS^+^ cells (Figures 3-4). When compared side-by-side, the reconstitution potential of BM LK201^+^FLT3^−^CD48^−^CD150^+^ LT-HSC versus LKS^+^FLT3^−^CD48^−^CD150^+^LT-HSC when sorted from LPS-treated mice (Figure 5) were similar and capable of multi-linage long-term reconstitution with similar RU content. However when BM HSPC from LPS-treated mice are sorted into either LK201^+^ or LKS^+^ phenotype then transplanted, the LK201^+^ sorted cells demonstrate a significant 4.4-fold higher reconstitution potential (Figure 3F) over sorted LKS^+^ phenotype cells. As both phenotypic LT-HSC populations contain the similar number of reconstituting HSC, the most likely explanation for this disparity is that the LKS^+^ compartment in LPS-treated mice becomes contaminated with CD201^−^ lineage-committed progenitor cells that have temporarily upregulated SCA1 in response to the LPS, but are devoid of long-term repopulation potential. For this reason, HSC phenotype gating strategies that incorporate CD201 based gating offers superior delineation of HSC, MPP and myeloid progenitor subsets in mice challenged with LPS.

A recent study claimed that LKS^+^FLT3^−^CD48^−^CD150^+^CD201^−^ cells contain myeloid biased HSC [13]. However in our hands, LK201^−^ cells from either saline-treated or LPS-treated mice gave very low levels of chimerism and reconstitution potential compared to LK201^+^ cells in a side-by-side comparison, suggesting that very few HSC are contained within CD201^−^populations.

We also found that due to strong SCA1 up-regulation on LK201^−^ progenitors in response to LPS, the numbers of myeloid progenitors in femurs of LPS-treated mice may have been previously overestimated when using the LKS^−^ gate compared to the LK201^−^ gate.

As LPS induces the release of type 1 and type 2 IFNs which are known to increase SCA1 expression on HSPC in the BM [6–9,37], it is reasonable to infer that similar observations would have been made in response to the administration of recombinant IFNs or pathogen-associated molecular patterns that elicit a strong IFN response.

An important observation is that SCA1 up-regulation in response to LPS was not limited to strains of the Ly6A.2 haplotype such as C57BL/6, but also occurred in the Ly6A.1 haplotype strain BALB/c which normally express very low levels of SCA1 in steady-state [3]. As CD201 is expressed by long-term reconstituting HSC in BALB/c and NOD strains [5,18], CD201 is likely to be a more suitable marker to identify HSC, MPP and HPC subsets in steady-state conditions or following inflammation and IFN-mediated signaling in a wide variety of mouse strains.

Having shown that CD201 expression better delineates reconstituting LT-HSC and HSPC subsets following LPS treatment, we next re-assessed the effect of LPS on HSC and MPP numbers (Figure 2 and Supplemental Figure 4) and proliferation (Figure 6) based on CD201 gating instead of SCA1 gating. LPS increased CD201^+^ LT-HSC, ST-HSC, MPP3 and MPP4 proliferation in line with previous studies showing an increase in HSC and MPP proliferation based on SCA1 positivity [12,14,15,38]. However, because of the strong SCA1 upregulation on LT-HSC and MPPs, the increase in the number of SCA1^+^ MPP2 and MPP3 per femur in response to LPS was overestimated compared to CD201^+^ MPP2 and MPP3 (Figure 2K,L and Supplementary Table 2), while the reduction in myeloid progenitor numbers following LPS treatment was also overestimated when using the LKS^−^ gate instead of LK201^−^ gate (Supplemental Table 3).

In respect to TLR-4 downstream signaling pathways involved in LPS-mediated SCA1 up-regulation on HSPC, our study shows that it was dependent on both TRIF and MYD88 mediated signaling in contrast to a previous study where SCA1 upregulation in response to LPS was reported to be only TRIF-dependent [14]. These different results may be explained by the fact that we analyzed SCA1 expression on LK cells whereas Lin^−^ cells were analyzed in the other study. As a proportion of quiescent BM Lin^−^ KIT^−^ cells express SCA1 but do not contain any hematopoietic activity [39], there presence may partly explain the difference in findings between these two studies.

Finally, we re-assessed HSPC mobilization in response to LPS based on LK201^+^ gating as we were concerned that SCA1 upregulation may lead to an overestimation of HSC and MPP subsets in the blood. Indeed, we found that the number of mobilized LKS^+^ cells was 3.6-fold higher than mobilized LK201^+^ cells however mobilized phenotypic LT-HSC were similar when using either CD201 or SCA1 for the gating. Likewise, we found that LPS-mediated HSPC mobilization into the blood also required expression of both MYD88 and TRIF as deletion of either of these 2 genes was sufficient to prevent HSPC mobilization. In view of this result, it was surprising that LPS-mediated CXCL12 protein and *Vcam1*-transcript down-regulation in BM was MYD88-dependent only. This suggests that additional mechanisms to CXCL12 and VCAM1 downregulation are involved in LPS-induced HSPC mobilization downstream of TRIF.

Several alternative mouse HSC markers have been proposed. Expression of the mCherry fluorescent reported knocked-in the *Fdg5* gene identifies HSC in steady-state [40] and inflammatory conditions following poly(I:C) [10] or IFN-γ [41] and its expression in HSC remains insensitive to these inflammatory stimuli. However, there may be some barriers to scientists in obtaining these genetically engineered mice. CD86 has also been proposed to be a more reliable marker of mouse HSC than SCA1 in inflammatory conditions [11]. Although there is good evidence that CD86 is expressed on reconstituting mouse HSC [11], we could not find evidence from the literature that it is expressed by human reconstituting HSC. Finally endomucin [42–44] and endothelial cell-specific adhesion molecule (ESAM) [45] are also markers of human and mouse LT-HSC. However, whether the expression of these cell surface antigens is altered following challenge with LPS or IFNs has not yet been reported.

In conclusion, our results show that unlike SCA1, the expression of CD201 is not dramatically upregulated on BM HSPC during broad inflammation triggered by LPS. Thus CD201-based analysis better identifies HSCs during inflammation. We therefore propose to use CD201 instead of SCA1 to identify HSPC subsets in the mouse because unlike SCA1, CD201 is *1)* expressed on HSPC in a wider variety of mouse strains (such as NOD and Ly6A.1 mouse strains [5,17,18], *2)* CD201 is also expressed on primate and human HSC [20,46,47] and *3)* CD201 expression is less affected following inflammatory challenges such as LPS and IFNs as demonstrated herein. Of note, it has recently been shown that SCA1 also becomes upregulated in leptin receptor positive BM mesenchymal stromal cells in response to poly(I:C) which induces a strong IFN response [48] suggesting that SCA1 upregulation is a marker of activated IFN-mediated signaling in both HSPC and mesenchymal cells. Therefore, the use of CD201 could enable more meaningful comparisons of phenotypically defined CD201^+^ HSC between species.

## Acknowledgments

This work was supported by funds from the Mater Foundation. K.B. was supported by an American Society of Hematology Global Research Award and Mater Research Future Leaders Fellowship. We thank Translational Research Institute Flow cytometry core facility and biological resources facility for flow cytometry and animal experiments.

## Authorship

### Contribution

K.B. coordinated the work, planned, and performed experiments, interpreted results, wrote original draft, made the figures and edited the manuscript. S.S. and V.B. performed experiments. I.G.W. interpreted the results, and edited the manuscript. J.P.L. conceived the work, planned, and performed experiments, interpreted results, wrote and edited the manuscript.

## REFERENCES

1. Spangrude GJ, Heimfeld S, Weissman IL. Purification and characterization of mouse hematopoietic stem cells. Science. 1988;241:58–62. 10.1126/science.2898810.

2. Pietras EM, Reynaud D, Kang Y-A, et al. Functionally distinct subsets of lineage-biased multipotent progenitors control blood production in normal and regenerative conditions. Cell Stem Cell. 2015;17:35–46. 10.1016/j.stem.2015.05.003.

3. Spangrude G, Brooks D. Mouse strain variability in the expression of the hematopoietic stem cell antigen Ly-6A/E by bone marrow cells. Blood. 1993;82:3327–32. 10.1182/blood.V82.11.3327.3327.

4. Chilton PM, Rezzoug F, Ratajczak MZ, et al. Hematopoietic stem cells from NOD mice exhibit autonomous behavior and a competitive advantage in allogeneic recipients. Blood. 2005;105:2189–97. 10.1182/blood-2004-07-2757.

5. Nowlan B, Williams ED, Doran MR, Levesque J-P. CD27, CD201, FLT3, CD48, and CD150 cell surface staining identifies long-term mouse hematopoietic stem cells in immunodeficient non-obese diabetic severe combined immune deficient-derived strains. Haematologica. 2020;105:71–82. 10.3324/haematol.2018.212910.

6. Essers MAG, Offner S, Blanco-Bose WE, et al. IFNα activates dormant haematopoietic stem cells in vivo. Nature. 2009;458:904–08. 10.1038/nature07815.

7. Pietras EM, Lakshminarasimhan R, Techner J-M, et al. Re-entry into quiescence protects hematopoietic stem cells from the killing effect of chronic exposure to type I interferons. J Exp Med. 2014;211:245–62. 10.1084/jem.20131043.

8. Chen J, Feng X, Desierto MJ, Keyvanfar K, Young NS. IFN-γ-mediated hematopoietic cell destruction in murine models of immune-mediated bone marrow failure. Blood. 2015;126:2621–31. 10.1182/blood-2015-06-652453.

9. Umemoto T, Matsuzaki Y, Shiratsuchi Y, et al. Integrin αvβ3 enhances the suppressive effect of interferon-γ on hematopoietic stem cells. EMBO J. 2017;36:2390–403. 10.15252/embj.201796771.

10. Bujanover N, Goldstein O, Greenshpan Y, et al. Identification of immune-activated hematopoietic stem cells. Leukemia. 2018;32:2016–20. 10.1038/s41375-018-0220-z.

11. Kanayama M, Izumi Y, Yamauchi Y, et al. CD86-based analysis enables observation of bona fide hematopoietic responses. Blood. 2020;136:1144–54. 10.1182/blood.2020004923.

12. Takizawa H, Fritsch K, Kovtonyuk LV, et al. Pathogen-induced TLR4-TRIF innate immune signaling in hematopoietic stem cells promotes proliferation but reduces competitive fitness. Cell Stem Cell. 2017;21:225–40.e5. 10.1016/j.stem.2017.06.013.

13. Vanickova K, Milosevic M, Ribeiro Bas I, et al. Hematopoietic stem cells undergo a lymphoid to myeloid switch in early stages of emergency granulopoiesis. EMBO J. 2023;42:e113527. 10.15252/embj.2023113527.

14. Zhang H, Rodriguez S, Wang L, et al. Sepsis induces hematopoietic stem cell exhaustion and myelosuppression through distinct contributions of TRIF and MYD88. Stem Cell Rep. 2016;6:940–56. 10.1016/j.stemcr.2016.05.002.

15. Demel UM, Lutz R, Sujer S, et al. A complex proinflammatory cascade mediates the activation of HSCs upon LPS exposure in vivo. Blood Adv. 2022;6:3513–28. 10.1182/bloodadvances.2021006088.

16. Kosanovic S, Vanickova K, Milosevic M, et al. Distinct transcriptional changes in hematopoietic progenitor subsets on LPS-induced emergency granulopoiesis. Exp Hematol. 2025;147:104792. 10.1016/j.exphem.2025.104792.

17. Balazs AB, Fabian AJ, Esmon CT, Mulligan RC. Endothelial protein C receptor (CD201) explicitly identifies hematopoietic stem cells in murine bone marrow. Blood. 2006;107:2317–21. 10.1182/blood-2005-06-2249.

18. Vazquez SE, Inlay MA, Serwold T. CD201 and CD27 identify hematopoietic stem and progenitor cells across multiple murine strains independently of Kit and Sca-1. Exp Hematol. 2015;43:578–85. 10.1016/j.exphem.2015.04.001.

19. Vanuytsel K, Villacorta-Martin C, Lindstrom-Vautrin J, et al. Multi-modal profiling of human fetal liver hematopoietic stem cells reveals the molecular signature of engraftment. Nat Comm. 2022;13:1103. 10.1038/s41467-022-28616-x.

20. Anjos-Afonso F, Buettner F, Mian SA, et al. Single cell analyses identify a highly regenerative and homogenous human CD34+ hematopoietic stem cell population. Nat Comm. 2022;13:2048. 10.1038/s41467-022-29675-w.

21. Adachi O, Kawai T, Takeda K, et al. Targeted disruption of the MyD88 gene results in loss of IL-1- and IL-18-mediated function. Immunity. 1998;9:143–50. 10.1016/S1074-7613(00)80596-8.

22. Yamamoto M, Sato S, Hemmi H, et al. Role of adaptor TRIF in the MyD88-independent toll-like receptor signaling pathway. Science. 2003;301:640–43. 10.1126/science.1087262.

23. Levesque JP, Takamatsu Y, Nilsson SK, Haylock DN, Simmons PJ. Vascular cell adhesion molecule-1 (CD106) is cleaved by neutrophil proteases in the bone marrow following hematopoietic progenitor cell mobilization by granulocyte colony-stimulating factor. Blood. 2001;98:1289–97. 10.1182/blood.v98.5.1289.

24. Levesque JP, Hendy J, Takamatsu Y, Williams B, Winkler IG, Simmons PJ. Mobilization by either cyclophosphamide or granulocyte colony-stimulating factor transforms the bone marrow into a highly proteolytic environment. Exp Hematol. 2002;30:440–49. 10.1016/s0301-472x(02)00788-9.

25. Levesque JP, Hendy J, Takamatsu Y, Simmons PJ, Bendall LJ. Disruption of the CXCR4/CXCL12 chemotactic interaction during hematopoietic stem cell mobilization induced by GCSF or cyclophosphamide. J Clin Invest. 2003;111:187–96. 10.1172/JCI15994.

26. Winkler IG, Sims NA, Pettit AR, et al. Bone marrow macrophages maintain hematopoietic stem cell (HSC) niches and their depletion mobilizes HSCs. Blood. 2010;116:4815–28. 10.1182/blood-2009-11-253534.

27. Purton LE, Scadden DT. Limiting factors in murine hematopoietic stem cell assays Cell Stem Cell. 2007;1:262–70. 10.1016/j.stem.2007.08.016.

28. Winkler IG, Pettit AR, Raggatt LJ, et al. Hematopoietic stem cell mobilizing agents G-CSF, cyclophosphamide or AMD3100 have distinct mechanisms of action on bone marrow HSC niches and bone formation. Leukemia. 2012;26:1594–601. 10.1038/leu.2012.17

29. Barbier V, Nowlan B, Levesque JP, Winkler IG. Flow cytometry analysis of cell cycling and proliferation in mouse hematopoietic stem and progenitor cells. Methods Mol Biol. 2012;844:31–43. 10.1007/978-1-61779-527-5_3.

30. Morcos MNF, Li C, Munz CM, et al. Fate mapping of hematopoietic stem cells reveals two pathways of native thrombopoiesis. Nat Commun. 2022;13:4504. 10.1038/s41467-022-31914-z.

31. Wilson A, Laurenti E, Oser G, et al. Hematopoietic stem cells reversibly switch from dormancy to self-renewal during homeostasis and repair. Cell. 2008;135:1118–29. 10.1016/j.cell.2008.10.048.

32. Forristal CE, Winkler IG, Nowlan B, Barbier V, Walkinshaw G, Levesque J-P. Pharmacologic stabilization of HIF-1α increases hematopoietic stem cell quiescence in vivo and accelerates blood recovery after severe irradiation. Blood. 2013;121:759–69. 10.1182/blood-2012-02-408419.

33. Burberry A, Zeng Melody Y, Ding L, et al. Infection mobilizes hematopoietic stem cells through cooperative NOD-like receptor and toll-like receptor signaling. Cell Host Microbe. 2014;15:779–91. 10.1016/j.chom.2014.05.004.

34. Bisht K, Tay J, Wellburn RN, et al. Bacterial lipopolysaccharides suppress erythroblastic islands and erythropoiesis in the bone marrow in an extrinsic and G-CSF-, IL-1-, and TNF-independent manner. Front Immunol. 2020;11:2548. 10.3389/fimmu.2020.583550.

35. Foudi A, Jarrier P, Zhang Y, et al. Reduced retention of radioprotective hematopoietic cells within the bone marrow microenvironment in CXCR4-/-chimeric mice. Blood. 2006;107:2243–51. 10.1182/blood-2005-02-0581.

36. Craddock CF, Nakamoto B, Andrews RG, Priestley GV, Papayannopoulou T. Antibodies to VLA4 integrin mobilize long-term repopulating cells and augment cytokine-induced mobilization in primates and mice. Blood. 1997;90:4779–88. 10.1182/blood.V90.12.4779.

37. Demerdash Y, Kain B, Essers MAG, King KY. Yin and Yang: The dual effects of interferons on hematopoiesis. Exp Hematol. 2021;96:1–12. 10.1016/j.exphem.2021.02.002.

38. Esplin BL, Shimazu T, Welner RS, et al. Chronic exposure to a TLR ligand injures hematopoietic stem cells. J Immunol. 2011;186:5367–75. 10.4049/jimmunol.1003438.

39. Randall TD, Weissman IL. Characterization of a population of cells in the bone marrow that phenotypically mimics hematopoietic stem cells: resting stem cells or mystery population? STEM CELLS. 1998;16:38–48. 10.1002/stem.160038.

40. Gazit R, Mandal PK, Ebina W, et al. Fgd5 identifies hematopoietic stem cells in the murine bone marrow. J Exp Med. 2014;211:1315–31. 10.1084/jem.20130428.

41. Morales-Mantilla DE, King KY. FGD5 marks a subpopulation of HSPCs that resists IFN-γ-mediated differentiation. Exp Hematol. 2022;112-113:35-43. 10.1016/j.exphem.2022.06.001.

42. Engelhard S, Estruch M, Qin S, et al. Endomucin marks quiescent long-term multi-lineage repopulating hematopoietic stem cells and is essential for their transendothelial migration. Cell Rep. 2024;43:114475. 10.1016/j.celrep.2024.114475.

43. Matsubara A, Iwama A, Yamazaki S, et al. Endomucin, a CD34-like sialomucin, marks hematopoietic stem cells throughout development. J Exp Med. 2005;202:1483–92. 10.1084/jem.20051325.

44. Reckzeh K, Kizilkaya H, Helbo AS, et al. Human adult HSCs can be discriminated from lineage-committed HPCs by the expression of endomucin. Blood Adv. 2018;2:1628. 10.1182/bloodadvances.2018015743.

45. Ooi AGL, Karsunky H, Majeti R, et al. The adhesion molecule ESAM1 is a novel hematopoietic stem cell marker. STEM CELLS. 2009;27:653–61. 10.1634/stemcells.2008-0824.

46. Fares I, Chagraoui J, Lehnertz B, et al. EPCR expression marks UM171-expanded CD34+ cord blood stem cells. Blood. 2017;129:3344–51. 10.1182/blood-2016-11-750729.

47. Papa L, Djedaini M, Martin TC, et al. Limited mitochondrial activity coupled with strong expression of CD34, CD90 and EPCR determines the functional fitness of ex vivo expanded human hematopoietic stem cells. Front Cell Dev Biol. 2020;8:92348. 10.3389/fcell.2020.592348.

48. Swann JW, Zhang R, Verovskaya EV, et al. Inflammation perturbs hematopoiesis by remodeling specific compartments of the bone marrow niche. Blood. 2025: in press. 10.1182/blood.2025029513.

