## Supplementary tables and figures for "Endothelial protein C receptor CD201 is a better marker than SCA1 to identify mouse long-term reconstituting hematopoietic stem cells following septic challenge"

Supplemental material.

Supplemental tables 1-3
Supplemental figures 1-10

**Supplemental Table 1.** Fluorescent antibodies and reagents used for flow cytometry.

| Reagents and resources | Suppliers | Catalog | Clone | Dilution |
|---|---|---|---|---|
| Anti-mouse CD3ε-Percpcy5.5 | BioLegend | 100328 | 145-2C11 | 1/200 |
| Anti-mouse CD5- Percpcy5.5 | BioLegend | 100624 | 53-7.3 | 1/200 |
| Anti-mouse CD45R (B220)- Percpcy5.5 | BioLegend | 103236 | RA3-6B2 | 1/200 |
| Anti-mouse/human CD11b- Percpcy5.5 | BioLegend | 101228 | M1/70 | 1/200 |
| Anti-mouse Gr1- Percpcy5.5 | BioLegend | 108428 | RB6-8C5 | 1/200 |
| Anti-mouse Ter119- Percpcy5.5 | BioLegend | 116228 | TER-119 | 1/200 |
| Anti-mouse CD3ε-FITC | BioLegend | 100306 | 145-2C11 | 1/200 |
| Anti-mouse CD5-FITC | BioLegend | 100606 | 53-7.3 | 1/200 |
| Anti-mouse/human CD45R (B220)-FITC | BioLegend | 103206 | RA3-6B2 | 1/200 |
| Anti-mouse/human CD11b-FITC | BioLegend | 101206 | M1/70 | 1/200 |
| Anti-mouse Gr1-FITC | BioLegend | 108406 | RB6-8C5 | 1/200 |
| Anti-mouse Ter119-FITC | BioLegend | 116206 | TER-119 | 1/200 |
| Anti-mouse CD117 (KIT)-APCCy7 | BioLegend | 105812 | 2B8 | 1/200 |
| Anti-mouse KIT-BV650 | BioLegend | 105839 | 2B8 | 1/200 |
| Anti-mouse SCA-1-FITC | BioLegend | 108105 | D7 | 1/200 |
| Anti-mouse SCA1-BV510 | BioLegend | 108129 | D7 | 1/100 |
| Anti-mouse SCA1-PECY7 | BioLegend | 108114 | D7 | 1/300 |
| Anti-mouse CD48-BV421 | BioLegend | 103427 | HM48-1 | 1/200 |
| Anti-mouse CD48-A700 | BioLegend | 103426 | HM48-1 | 1/200 |
| Anti-mouse CD150-PE | BioLegend | 115904 | TC15-12F12.2 | 1/200 |
| Anti-mouse CD150-PECy7 | BioLegend | 115914 | TC15-12F12.2 | 1/200 |
| Anti-mouse CD45-BV785 | BioLegend | 103149 | 30-F11 | 1/200 |
| Anti-mouse CD135 (FLT3)-PECF594 | BD Biosciences | 562537 | A2F10.1 | 1/150 |
| Anti-mouse CD201-Biotin | Biolegend | Cat#141508 | clone RCR-16 | 1/200 |
| Anti-mouse CD105-PE | BioLegend | 120408 | MJ 7/18 | 1/200 |
| Anti-mouse CD16/32-Percpcy5.5 | BioLegend | 101324 | 93 | 1/100 |
| Anti-mouse CD41-APCCy7 | BioLegend | 133928 | MWReg30 | 1/200 |
| Anti-mouse CD45.1-PE | BioLegend | 110708 | A20 | 1/200 |
| Anti-mouse CD45.2-APC | BioLegend | 109814 | 104 | 1/200 |
| Anti-mouse CD3ε -BV421 | BioLegend | 100341 | 145-2C11 | 1/200 |
| Anti-mouse/human CD11b-BV510 | BioLegend | 101245 | M1/70 | 1/200 |
| Anti-mouse LY6G-PECY7 | BioLegend | 127618 | 1A8 | 1/200 |
| Anti-mouse/human CD45R (B220)-APCCY7 | BioLegend | 03224 | RA3-6B2 | 1/200 |
| Streptavidin-APC | BioLegend | 405207 | | 1/200 |
| Anti-human Ki67-FITC | BD Biosciences | 51-36524X | | 1/100 |
| FC block (neutralizing mouse CD16/32) | Dr Louise Purton | | 2.4G2 | 50% |
| Hoechst33342 | ThermoFisher Scientific | 62249 | | 1/800 |
| FVS700 | BD Biosciences | 564997 | | 1/10,000 |

**Supplemental Table 2.** Percentage change of HSC and MPP numbers in the BM following in vivo LPS treatment compared to saline treatment

| **Populations** | **LK201+** | **LKS+** |
|---|---|---|
| LK | -53.8 ± 22.7 | 10.9 ± 56.7 |
| LT-HSC | -31.4 ± 48.1 | -47.2 ± 19.6 |
| ST-HSC | -84.5 ± 17.1 | -81.6 ± 26.5 |
| MPP2 | 220.0 ± 128.9 | 325.8 ± 202.7 |
| MPP3 | -34.9 ± 31.4 | 152.4 ± 142.3 |
| MPP4 | -68.3 ± 30.1 | -41.8 ± 30.1 |

**Supplemental Table 3.** Percentage of change of myeloid progenitor numbers in the BM following in vivo LPS treatment compared to saline treatment

| **Populations** | **LK201-** | **LKS-** |
|---|---|---|
| CMP | -60.9 ± 23.7 | -88.3 ± 9.6 |
| GMP | -67.0 ± 20.4 | -65.9 ± 23.8 |
| MEP | -72.8 ± 13.5 | -81.3 ± 14.9 |
| MegP | -35.9 ± 15.6 | -53.0 ± 32.0 |
| BFU-E | -78.2 ± 11.0 | -92.5 ± 4.0 |
| CFU-E | -77.4 ± 14.0 | -82.0 ± 14.5 |

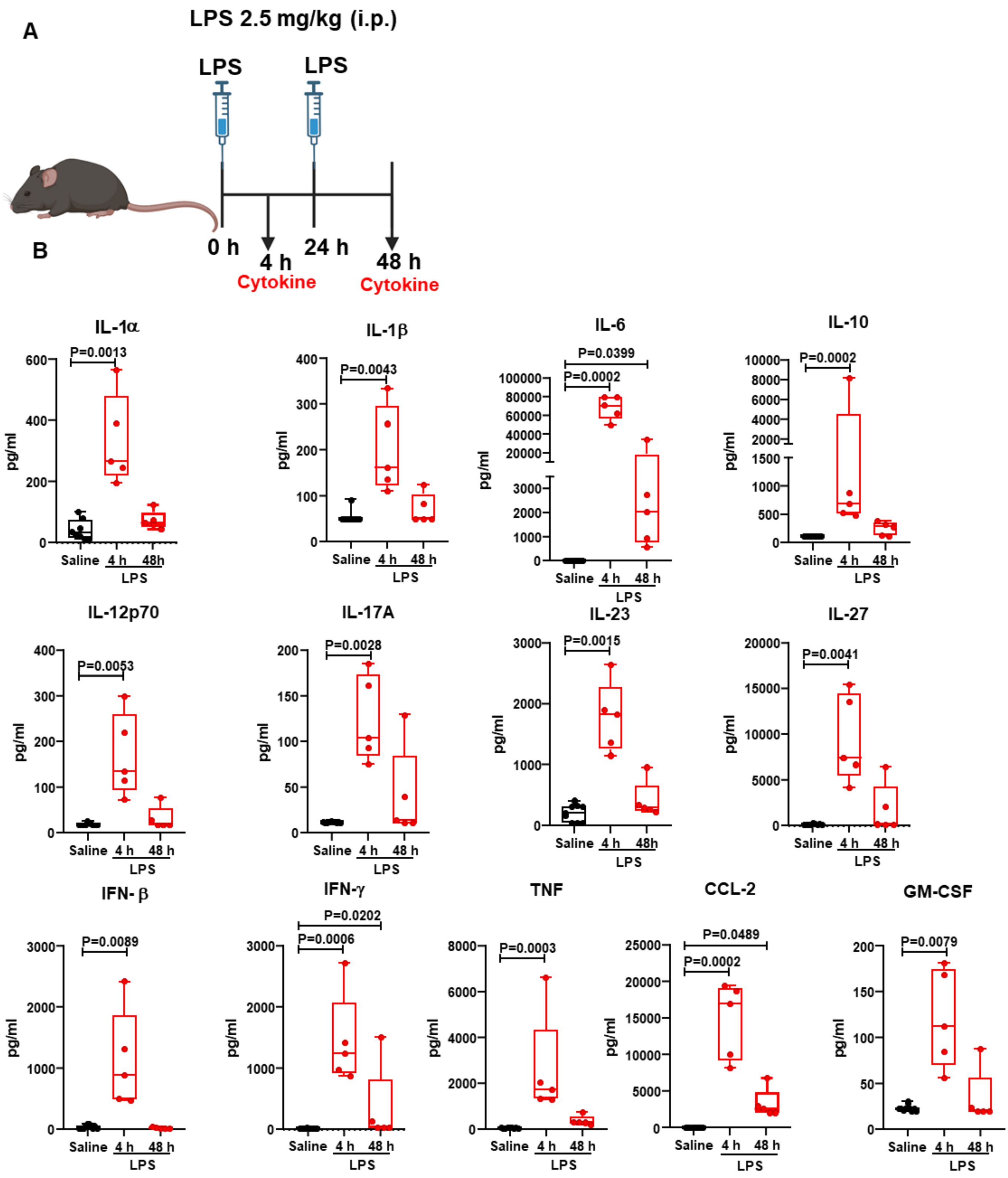


**Supplemental Figure 1.** LPS treatment increases inflammatory cytokines in mouse blood. (**A**) C57BL/6 mice were administered saline or LPS once daily for two consecutive days. Blood was collected either 4 or 48 hour after the last LPS injection for cytokine measurement. The figure was created with BioRender.com. (**B**) Cytokines concentrations were measured using Legendplex mouse inflammation panel (13 plex). Each dot represents a separate mouse. Bars are mean ± SD. Statistical significance was determined using non-parametric Kruskal Wallis multiple comparison test.

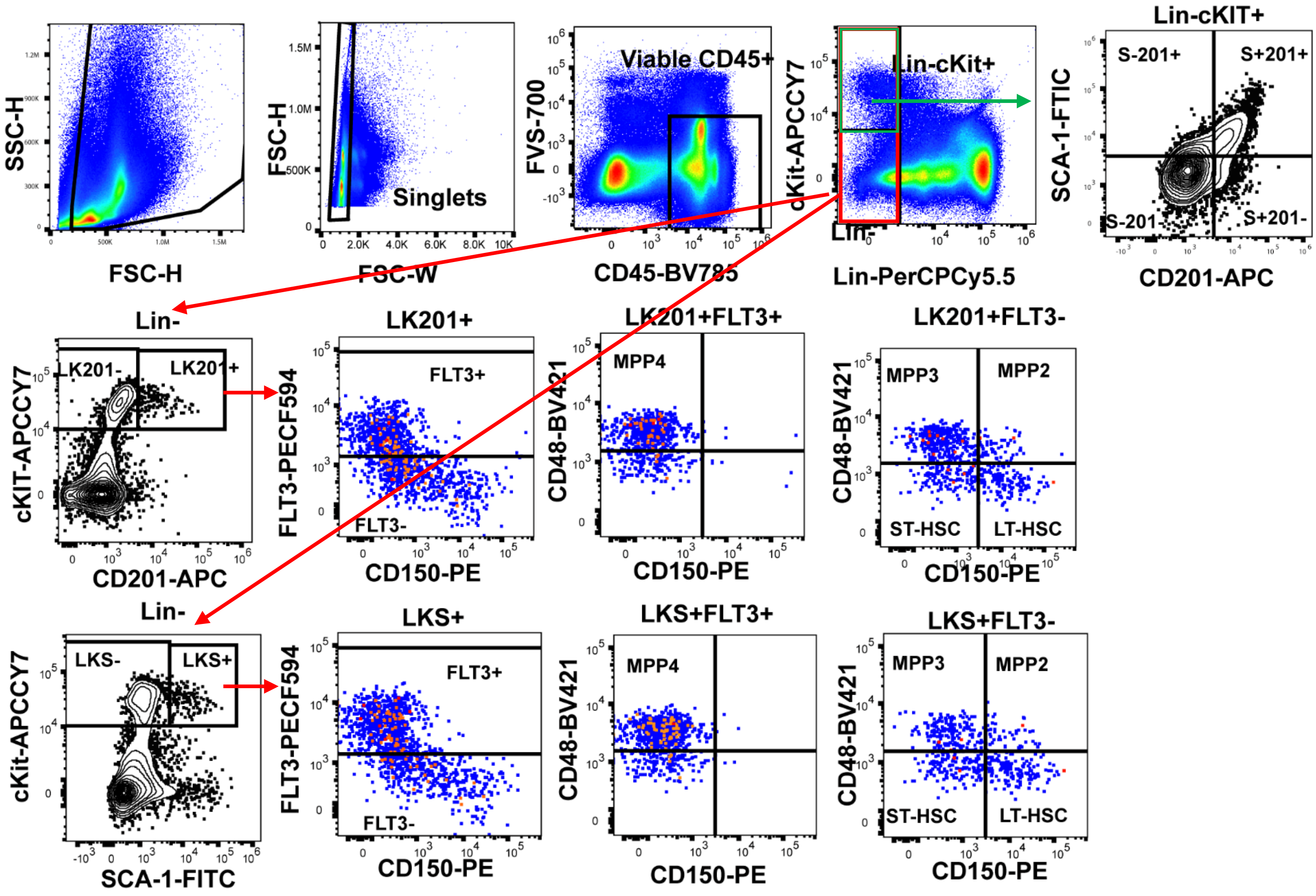


**Supplemental Figure 2.** Gating strategy to identify Lin-, LK, LK201$^{-/+}$, LKS$^{-/+}$, HSC and MPP subsets. This example is taken from a C57BL/6 mouse treated with saline for 48 hours. Singlet cells were gated according to forward and side scatter and then gated for viable (FVS700-negative) CD45$^{+}$ leukocytes. HSPCs were then identified as lineage-negative KIT$^{+}$ CD201$^{+}$ or SCA-1$^{+}$ and further gated as MPP4 (FLT3$^{+}$CD150$^{-}$CD48$^{+}$), MPP3 (FLT3$^{-}$CD150$^{-}$CD48$^{+}$), MPP2 (FLT3$^{-}$CD150$^{+}$CD48$^{+}$), ST-HSCs (FLT3$^{-}$CD150$^{-}$CD48$^{-}$) and LT-HSCs (FLT3$^{-}$CD150$^{+}$CD48$^{-}$).

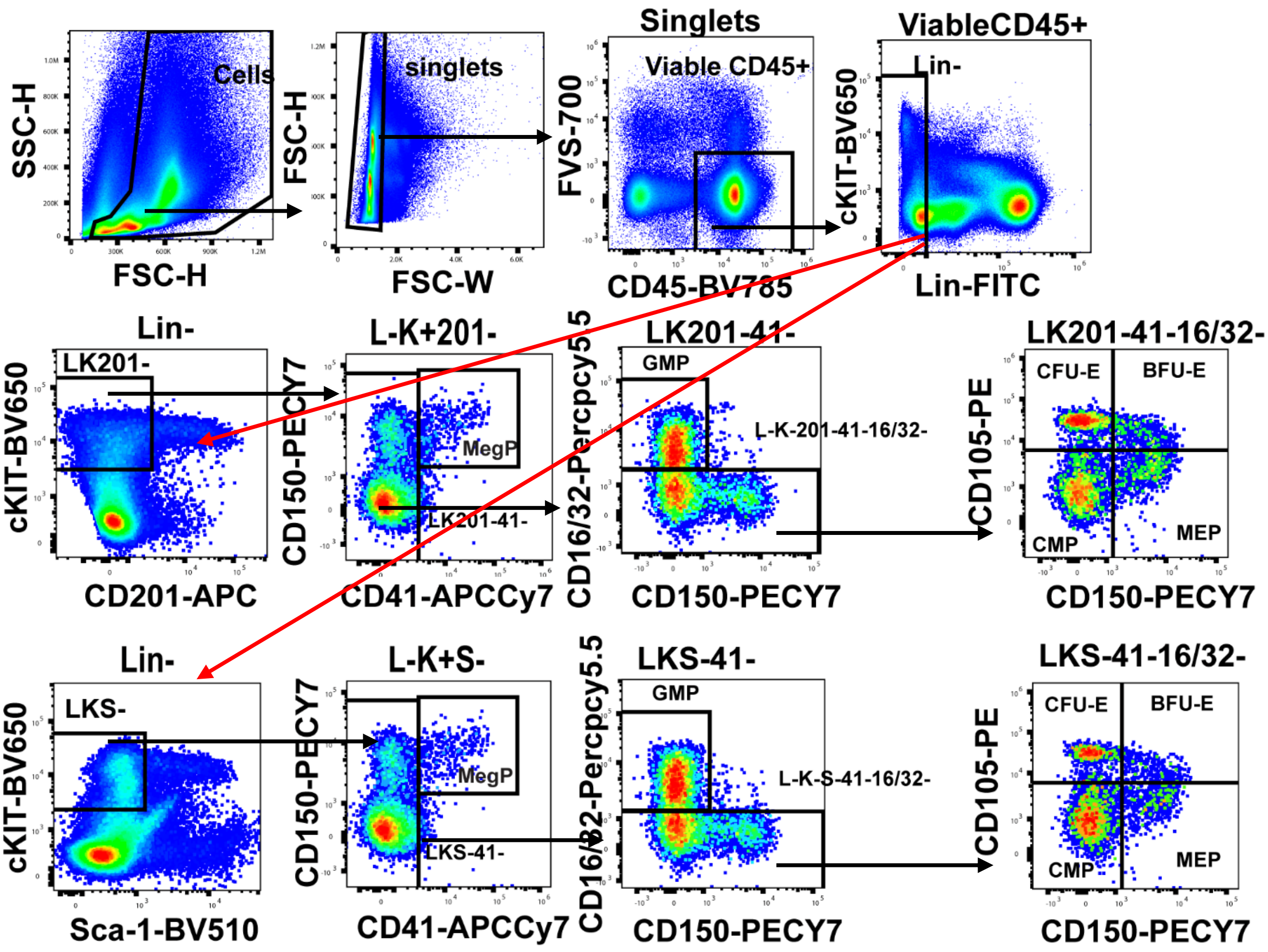


**Supplemental Figure 3**. Gating strategy to identify myeloid and erythroid progenitors. Example taken from a C57BL/6 mouse treated with saline. Singlet cells were gated according to forward and side scatter to then gated for viable FVS700$^{-}$ CD45$^{+}$ leukocytes followed by lineage-negative (Lin$^{-}$) cells. Lin$^{-}$KIT$^{+}$CD201$^{-}$ (LK201$^{-}$) or Lin$^{-}$KIT$^{+}$SCA1$^{-}$ (LKS-) cells were further gated as megakaryocyte progenitors MegP (CD41$^{+}$), granulocyte myeloid progenitors GMP (CD41$^{-}$CD16/32$^{+}$), common myeloid progenitors CMP (CD41$^{-}$CD105$^{-}$CD150$^{-}$), myeloid erythroid progenitors MEP (CD41$^{-}$ CD105$^{-}$ CD150$^{+}$), burst forming unit–E BFU-E (CD41$^{-}$ CD105$^{+}$CD150$^{+}$). and colony forming unit-E CFU-E (CD41$^{-}$ CD105$^{+}$CD150$^{-}$).

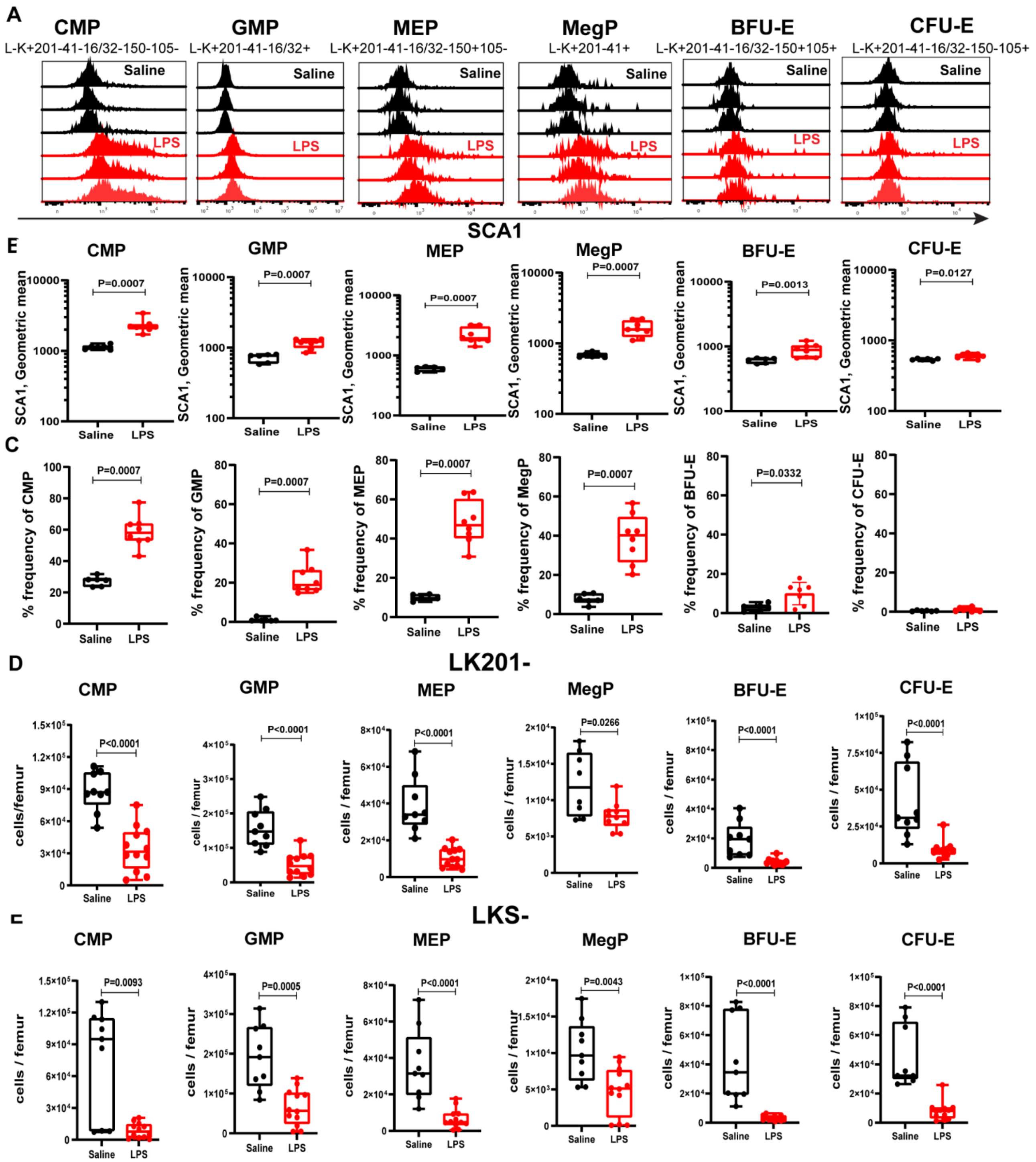


**Supplemental Figure 4.** LPS upregulates SCA1 expression on bone marrow myeloid and erythroid progenitors. C57BL/6 mice were injected twice with LPS once daily and then rested for 48 hours as in Figure 1A. (**A**) SCA1 flow cytometry histograms, (**B**) SCA1 geometric mean fluorescence intensities and (**C**) percentage SCA1$^+$ cells in LK201$^-$CD41$^-$CD16/32$^-$CD150$^-$CD105$^-$ CMP, LK201$^-$CD41$^-$CD16/32$^+$CD150$^-$ GMP, LK201$^-$CD41$^-$CD16/32$^-$CD150$^+$CD105$^-$ MEP, LK201$^-$CD41$^+$CD150$^+$ MegP, LK201$^-$CD41$^-$CD16/32$^-$CD150$^+$CD105$^+$ BFU-E and LK201$^-$CD41$^-$CD16/32$^-$CD150$^-$CD105$^+$ CFU-E. (**D-E**) Quantification of CMP, GMP, MEP, MegP, BFU-E and CFU-E per femur based on LK201$^-$ (**D**) or LKS$^-$ gating (**E**).

In (**A**) SCA1 histograms for HPC from 3 representative saline-treated (black histograms) and LPS-treated (red histograms) mice. In (**B-E**), each dot represents a separate mouse. Statistical significance was calculated using a non-parametric Mann-Whitney test.

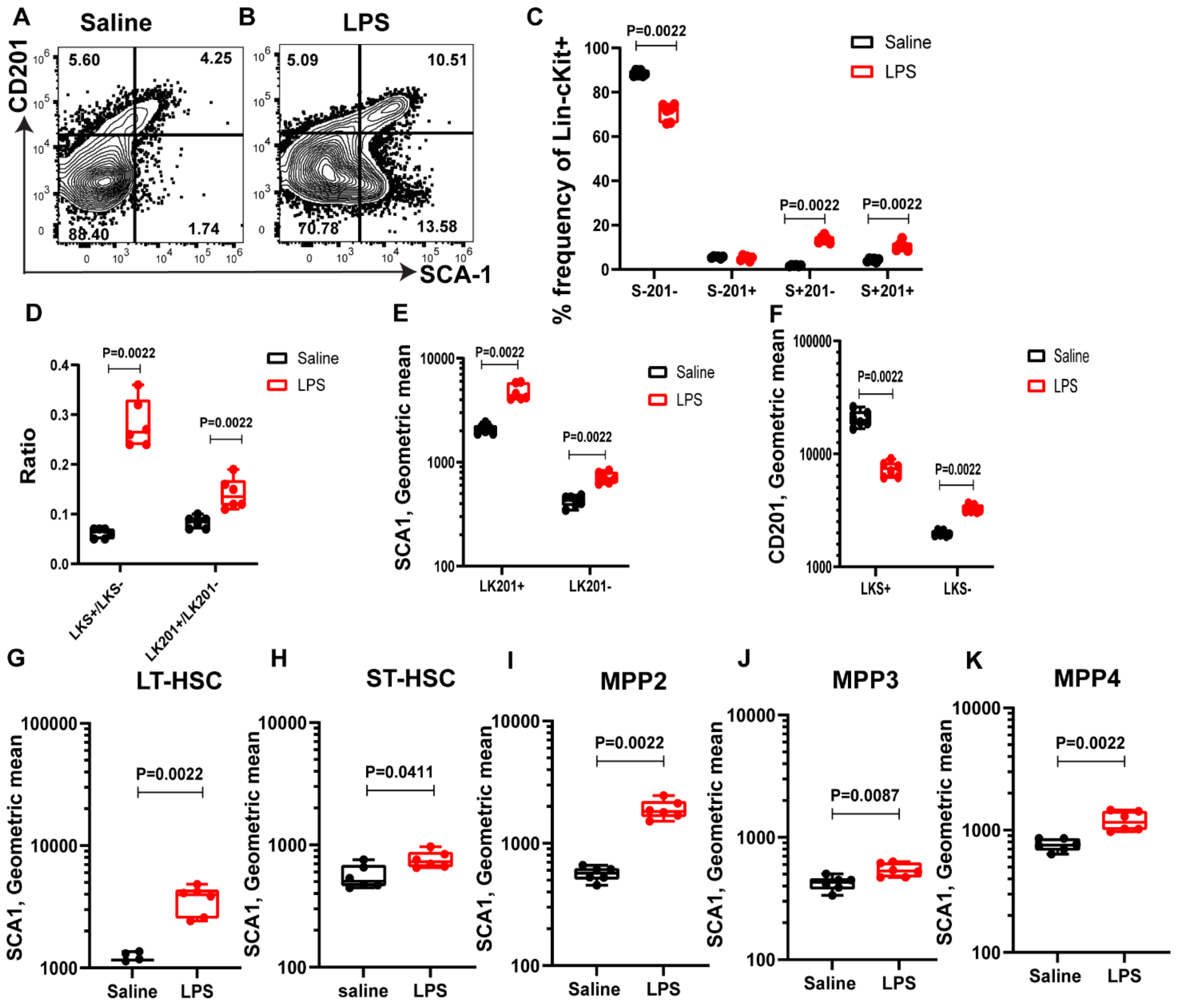


**Supplemental Figure 5.** LPS upregulates SCA1 expression on bone marrow HSC and MPP population in BALB/c mice. BALB/c mice were injected twice with LPS once daily and then rested for 48 hours as in Figure 1A. (**A-C**) Representative flow cytometry contour plots showing CD201 and SCA1 expression on BM Lin$^{-}$ KIT$^{+}$ HSPC, LT-HSC and ST-HSC cells following LPS or saline treatment. (**C**) Frequencies of SCA$^{-}$CD201$^{-}$ (S$^{-}$201$^{-}$), SCA1$^{-}$CD201$^{+}$ (S$^{-}$201$^{+}$), SCA1$^{+}$CD201$^{-}$ (S$^{+}$201$^{-}$), and SCA1$^{+}$CD201$^{+}$ (S$^{+}$201$^{+}$) cells on BM Lin$^{-}$KIT$^{+}$ cells. **(G-K**) SCA1 mean fluorescence intensity on (G) LK201+FLT3-CD48-CD150+ LT-HSC, (H) LK201+ FLT3-CD48-CD150- ST-HSC, (**I**) LK201+FLT3-CD48+CD150+ MPP2, (**J**) LK201+FLT3-CD48+CD150- MPP3 and (**K**) LK201+FLT3+CD48+CD150- MPP4 after LPS (red dots) or saline (black dots) treatment. Each dot is a separate mouse with mean ± SD. P values were calculated using Mann-Whitney test.

Donor: LK201+

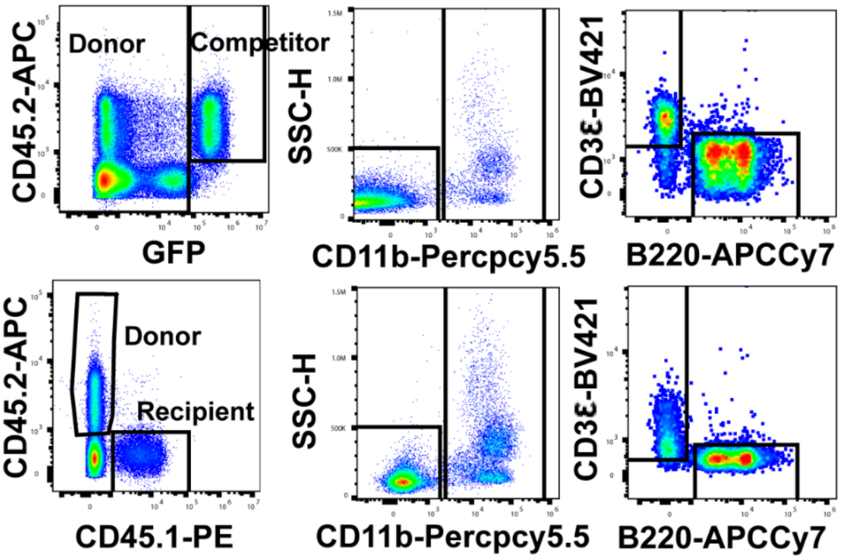


Donor: LK201-

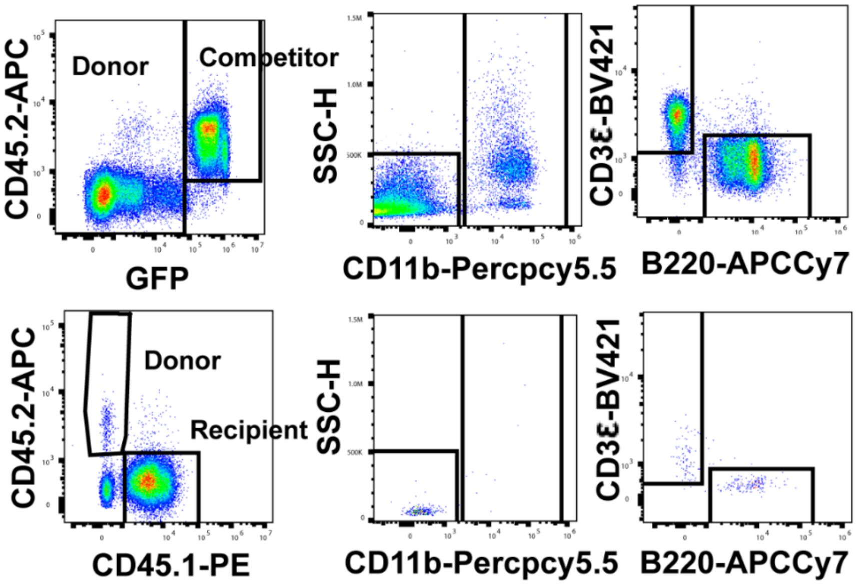


Donor: LKSca1+

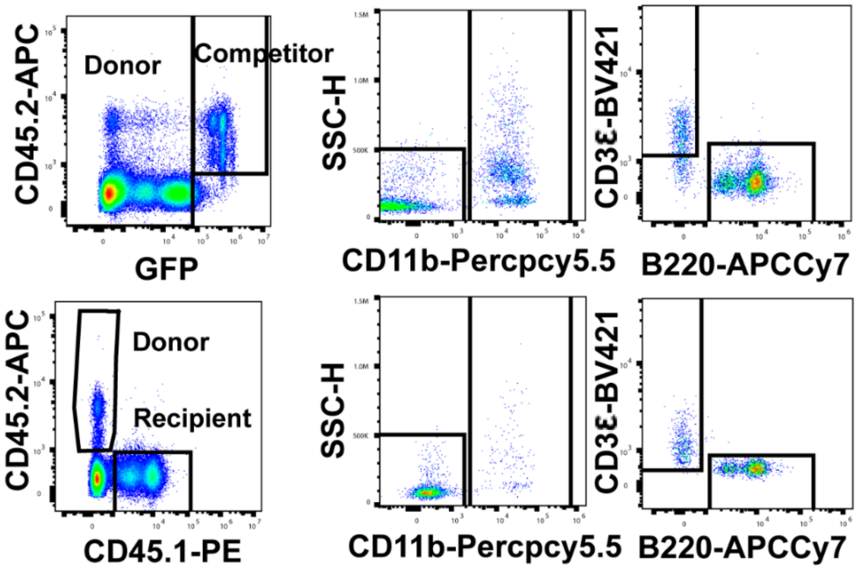

**Supplemental Figure 6**. Gating strategy to measure multilineage reconstitution from of LK201+, LK201- and LKS+ donor cells in long-term competitive repopulation assays. C57BL/6 mice were injected with saline or LPS for 48 hours and harvested after one day of recovery. $LK201^+$, $LK201^-$ and $LKS^+$ cells were sorted and transplanted into lethally irradiated B6.SJL CD45.1+ in competition with 400,000 BM cells from *Ubc*-GFP mice and mice were maintained for 20 weeks This example is taken from a B6.SJL CD45.1+ recipient mouse blood at 20 weeks post-transplantation. Singlet cells were gated according to forward and side scatter to then gated for viable $FVS700^-$ cells. followed by GFP- CD45.2+ (donor) and GFP+ (competitor) cells. Donor and competitor cells were then gated to numerate chimerism in CD11b+ myeloid cells, CD3+ T cells and B220+ B cells.

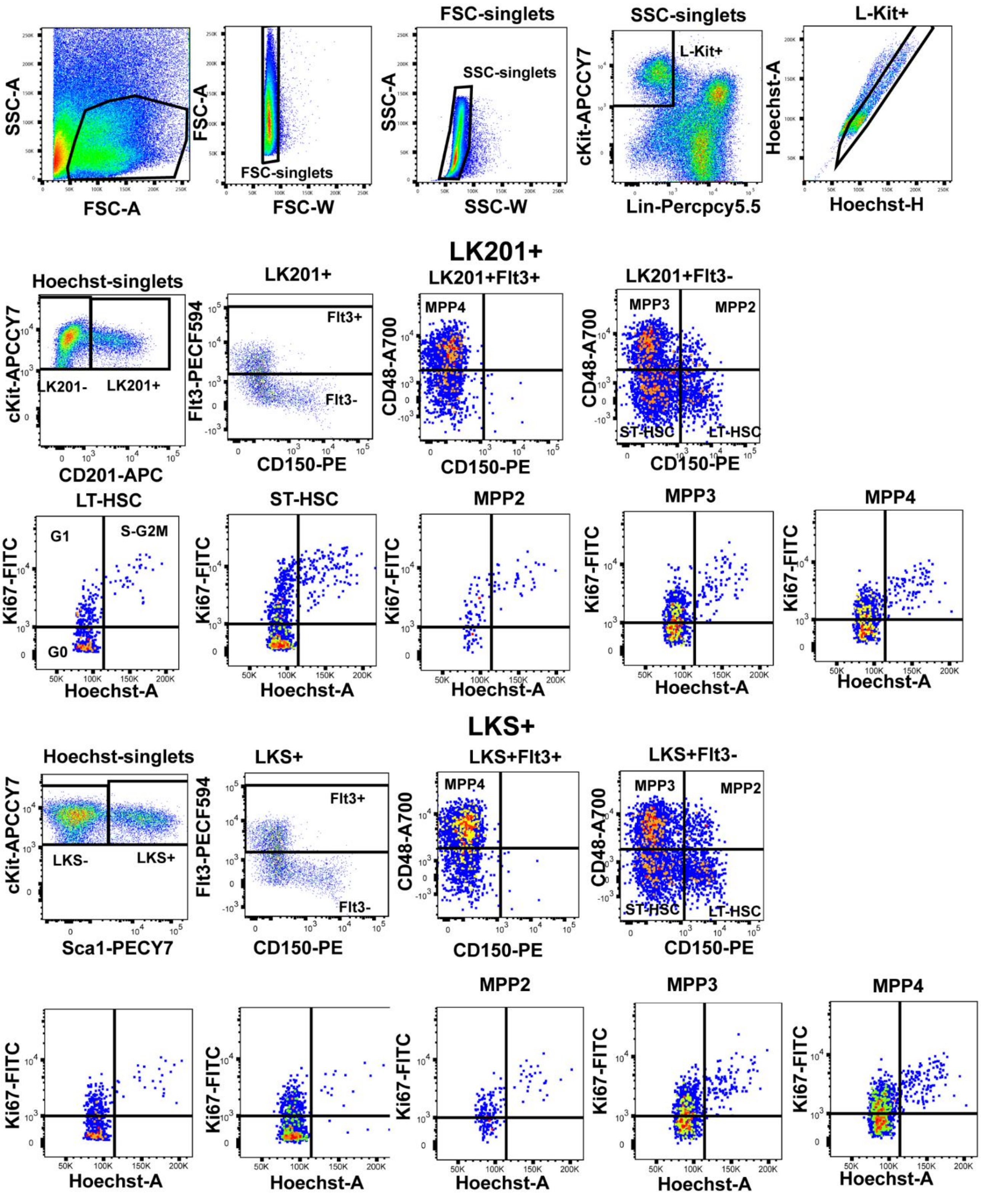


**Supplemental Figure 7.** Gating strategy for cell cycle analysis of BM HSPCs. This example is taken from a C57BL/6 mouse treated with saline. Singlet cells were gated according to forward and side scatter and then gated for lineage-negative ($Lin^-$) KIT+ cells. $Lin^-$ $KIT^+$ cells were further gated for singlets on Hoechst channel. Singlets were further gated as $Lin^-$ $KIT^+$ $CD201^+$ (LK201+) or $Lin^-$ $KIT^+$ $Sca1^+$ (LKS+). LK201+ and LKS+ cells were further gated as $FLT3^+CD150^-CD48^+$ for MPP4 , $FLT3^-CD150^-CD48+$ for MPP3, $FLT3^-CD150^+CD48^+$ for MPP2, $FLT3^-CD150^-CD48^-$ for ST-HSC and $FLT3^-CD150^+CD48^-$ for LT-HSC. Different phases of cell cycle were analyzed on MPPs and HSCs according to Ki67 staining and genomic DNA content (Hoechst).

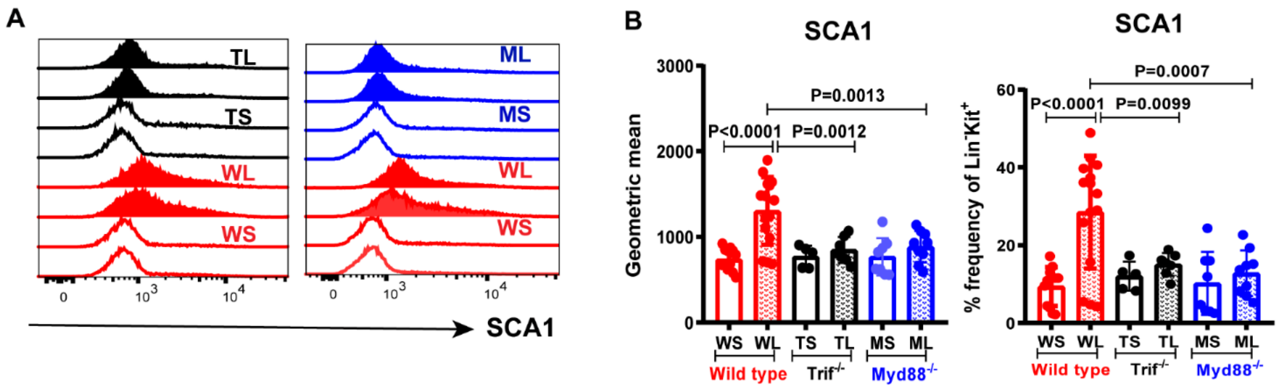


**Supplemental Figure 8**. LPS induces SCA1 expression via TRIF and MYD88 pathways in vivo. C57BL/6 mice were injected once daily with saline or LPS for 2 consecutive days and harvested after two days of recovery. (**A**) Representative flow cytometry histograms showing no increase in SCA1 expression in the BM after LPS in mice lacking either TRIF or MYD88. Each histogram represents SCA1 expression profile on LK cells from two separate WT mouse saline-treated (WS, red empty), WT LPS-treated (WL, red filled), TRIF KO saline-treated (TS, black empty), TRIF KO LPS-treated (TL, black filled), MyD88 KO saline-treated (MS, blue empty) or Myd88 KO LPS-treated (ML, blue filled) treated mice. (**B**) Geometric mean and frequency of SCA1 gated on LK cells from the same groups as in panel A. WS, WL, TS, TL, MS and ML groups denote mouse genotype and treatment as in Panel A. Each dot is a separate mouse with mean ± SD. Statistical significance was determined using One-way ANOVA with Sidak’s multiple comparison test.

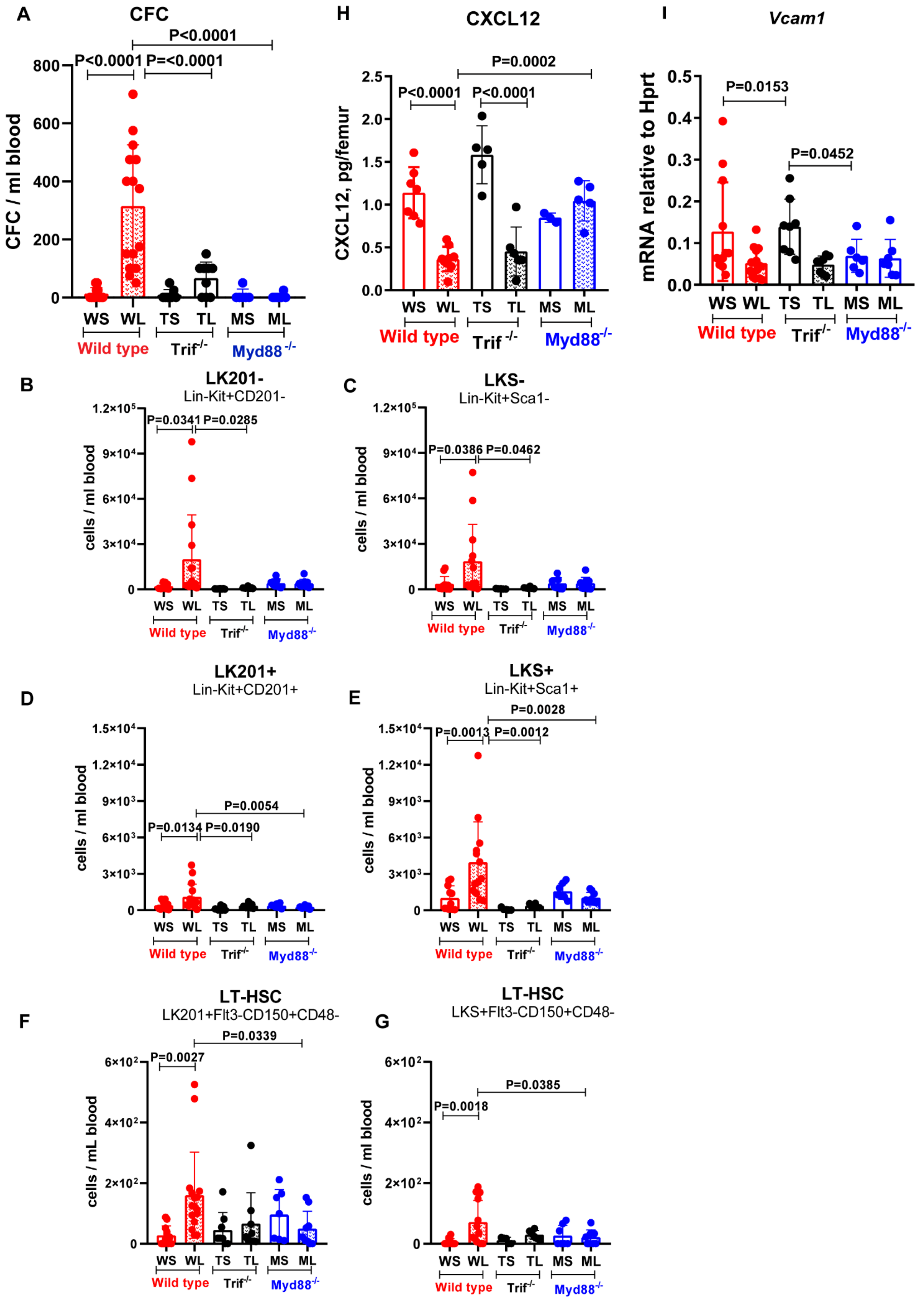
A
CFC
P<0.0001
P<0.0001
P=<0.0001
CFC / ml blood
0
200
400
600
800
WS WL TS TL MS ML
Wild type
Trif-/-
Myd88 -/-
H
CXCL12
P=0.0002
P<0.0001
P<0.0001
CXCL12, pg/femur
0.0
0.5
1.0
1.5
2.0
2.5
WS WL TS TL MS ML
Wild type
Trif -/-
Myd88-/-
I
Vcam1
P=0.0153
P=0.0452
mRNA relative to Hprt
0.0
0.1
0.2
0.3
0.4
0.5
WS WL TS TL MS ML
Wild type
Trif-/-
Myd88-/-
B
LK201-
Lin-Kit+CD201-
P=0.0341
P=0.0285
cells / ml blood
0
3×10⁴
6×10⁴
9×10⁴
1.2×10⁵
WS WL TS TL MS ML
Wild type
Trif-/-
Myd88-/-
C
LKS-
Lin-Kit+Sca1-
P=0.0386
P=0.0462
cells / ml blood
0
3×10⁴
6×10⁴
9×10⁴
1.2×10⁵
WS WL TS TL MS ML
Wild type
Trif-/-
Myd88-/-
D
LK201+
Lin-Kit+CD201+
P=0.0054
P=0.0134
P=0.0190
cells / ml blood
0
3×10³
6×10³
9×10³
1.2×10⁴
1.5×10⁴
WS WL TS TL MS ML
Wild type
Trif-/-
Myd88-/-
E
LKS+
Lin-Kit+Sca1+
P=0.0028
P=0.0013
P=0.0012
cells / ml blood
0
3×10³
6×10³
9×10³
1.2×10⁴
1.5×10⁴
WS WL TS TL MS ML
Wild type
Trif-/-
Myd88-/-
F
LT-HSC
LK201+Flt3-CD150+CD48-
P=0.0339
P=0.0027
cells / mL blood
0
2×10²
4×10²
6×10²
WS WL TS TL MS ML
Wild type
Trif-/-
Myd88-/-
G
LT-HSC
LKS+Flt3-CD150+CD48-
P=0.0385
P=0.0018
cells / ml blood
0
2×10²
4×10²
6×10²
WS WL TS TL MS ML
Wild type
Trif-/-
Myd88-/-

Figure 7

**Supplemental Figure 9.** LPS treatment increases HSPC mobilization into blood via TRIF and MYD88. C57BL/6 mice were injected once daily with saline or LPS for 2 consecutive days and harvested after two days of recovery. Numbers of CFC (**A**), $LK201^-$ (**B**), $LKS^-$ (**C**), $LK201^+$ (**D**), $LKS^+$ (**E**) HSPC and LT-HSC based on $LK201^+$ gating (**F**) or $LKS^+$ gating (**G**) in peripheral blood after LPS or saline treatment in wild-type (W), $TRIF^{-/-}$ (T) or $Myd88^{-/-}$ (M) mice treated with saline (S) or LPS (L). (**H**) CXCL12 concentration in BM fluids from the above treated mice. (**I**) qRT-PCR quantification of *Vcam1* mRNA on endosteal BM cells isolated from the above treated mice. Each dot is a separate mouse with mean ± SD. P-values were calculated using One-Way ANOVA with Sidak's multiple comparison test.

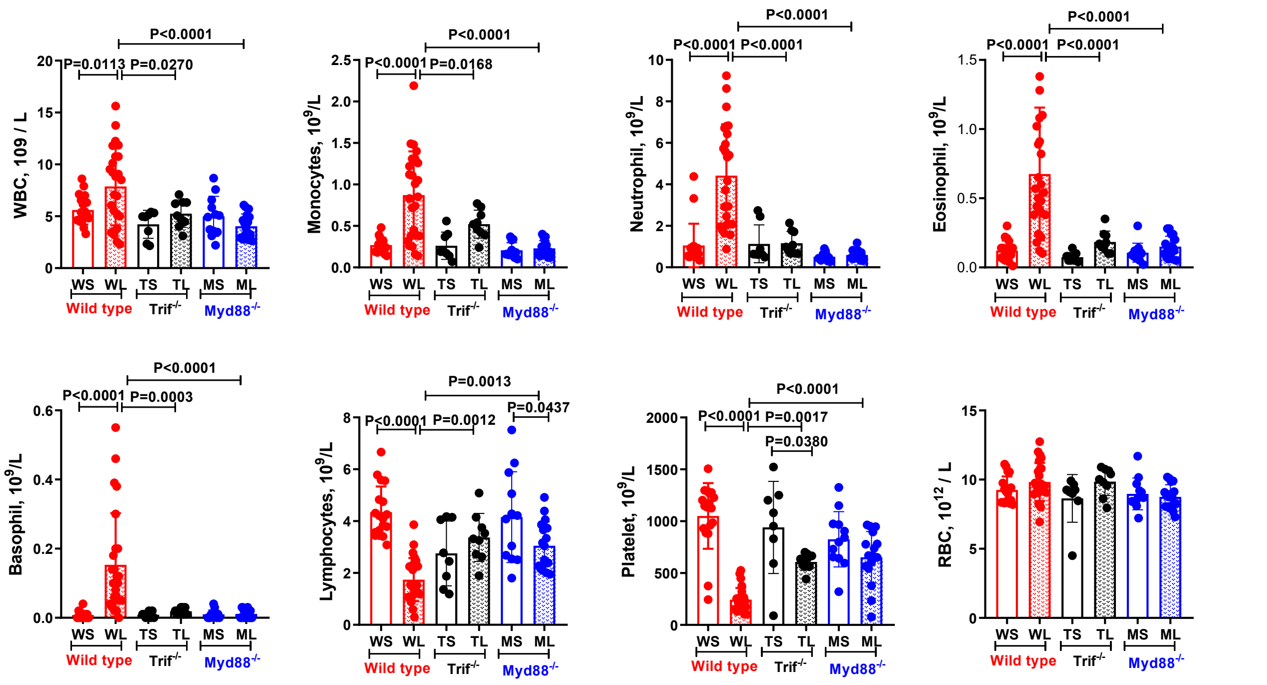


**Supplemental Figure 10.** Changes in blood cell numbers in response to LPS in wildtype, Myd88 or Trif knock-out mice. The number of white blood cells (WBC), monocytes, neutrophils, eosinophils, basophils, lymphocytes, platelets and red blood cells (RBC) were measured with a Mindray hematology analyzer. Data are from 4 pooled experiments performed few months intervals. Each dot is a separate mouse with mean ± SD. Statistical significance was determined using one way-ANOVA with Tukey's multiple comparison test.
